# HspB8 prevents aberrant phase transitions of FUS by chaperoning its folded RNA binding domain

**DOI:** 10.1101/2021.04.13.439588

**Authors:** Edgar E. Boczek, Julius Fürsch, Louise Jawerth, Marcus Jahnel, Marie Laura Niedermeier, Martine Ruer-Gruß, Kai-Michael Kammer, Laura Mediani, Jie Wang, Xiao Yan, Andrej Pozniakovski, Ina Poser, Daniel Mateju, Serena Carra, Simon Alberti, Anthony A. Hyman, Florian Stengel

## Abstract

Aberrant liquid-to-solid phase transitions of biomolecular condensates have been linked to various neurodegenerative diseases. However, the underlying molecular interactions that drive aging remain enigmatic. Here, we develop quantitative time-resolved crosslinking mass spectrometry to monitor protein interactions and dynamics inside condensates formed by the protein fused in sarcoma (FUS). We identify misfolding of the RNA recognition motif (RRM) of FUS as a key driver of condensate ageing. We demonstrate that the small heat shock protein HspB8 partitions into FUS condensates via its intrinsically disordered domain and prevents condensate hardening via condensate-specific interactions that are mediated by its α-crystallin domain (αCD). These αCD-mediated interactions are altered in a disease-associated mutant of HspB8, which abrogates the ability of HspB8 to prevent condensate hardening. We propose that stabilizing aggregation-prone folded RNA-binding domains inside condensates by molecular chaperones may be a general mechanism to prevent aberrant phase transitions.

## Introduction

Condensate formation by liquid-liquid phase separation (LLPS) leads to a local density change of proteins (1, 2). Liquid condensates can harden and evolve into less dynamic states with reduced fluidity and protein movement, leading to fibrillar assemblies that are often associated with disease (3–5). The molecular changes that underly these aberrant phase transitions, and the ways that cells prevent them, remain poorly understood.

Stress granules have been used as a model to study the role of phase separation in the formation of cellular condensates, as well as disease processes that arise from aberrant phase transitions. Stress granules are composed of RNA and translation factors (6–11). There is an increasing body of evidence that the ability of stress granule proteins to phase separate forms the basis for stress granule assembly (6–8). For instance, purified stress granule residing RNA-binding proteins (RBPs) phase separate into liquid droplets in vitro. Reconstituted droplets have physicochemical properties similar to stress granules in cells (4, 6–8). In vitro reconstituted molecular condensates are metastable and age into less dynamic amorphous drops, and fibrillar aggregates with time (4). These aggregates are reminiscent of protein aggregates seen in patients afflicted with age-related diseases such as Amyotrophic Lateral Sclerosis (ALS) and Frontotemporal Dementia (FTD). This process is referred to as molecular aging (4). Notably, the molecular aging process is accelerated by ALS-linked mutations in FUS and other RBPs (4, 5). This suggests that molecular aging of liquid condensates such as stress granules can be a disease process.

Recent studies have provided evidence for a link between stress granules and small heat shock proteins (sHSPs) (12–16). These ATP-independent chaperones hold unfolded proteins in a refolding-competent state and both their intrinsically disordered region (IDR) and folded alpha crystallin domain (αCD) contribute to this activity (17). sHSPs have been shown to accumulate in stress granules that undergo an aberrant conversion from a liquid to a solid-like state (12, 13). Aberrant stress granules have also been linked to disease and many sHSPs are associated with neurodegenerative disorders (18). This suggests that chaperones such as sHSPs may regulate the properties of stress granules and presumably also the molecular aging process of stress granule proteins such as FUS.

FUS has been a model protein to study both aberrant and physiological phase transitions (4, 19–24). It contains an intrinsically disordered prion-like low complexity domain (LCD) composed of only a small subset of amino acids, an RNA binding domain (RBD) containing intrinsically disordered RGG-rich motifs, a Zinc finger (ZnF) and a folded RNA recognition motif (RRM). It has been shown that phase separation of FUS family proteins is driven by multivalent interactions among tyrosine residues within its LCD and arginine residues of their RBDs (21, 22, 25). At least part of the LCD of FUS can assemble into amyloid fibrils (26) and the isolated FUS-LCD can adopt a hydrogel-like state that also depends on amyloid-like interactions (24). By contrast, the LCD appears to be disordered in liquid droplets, exhibiting no detectable secondary structure (27). Similar observations have been made for the LCD of hnRNPA2 (28). In addition to the LCD, the isolated FUS-RRM has been shown to spontaneously self-assemble into amyloid fibrils (29). However, the molecular mechanism by which proteins such as FUS undergo molecular aging are still unknown.

To determine the molecular changes during the aging of stress granule proteins such as FUS, it is critical to monitor protein-protein interactions (PPIs) and conformational dynamics within condensates. However, this has remained a major challenge. We and others showed previously that chemical crosslinking coupled to mass spectrometry (XL-MS) is well suited to map PPIs and that relative changes in crosslinking as probed by quantitative XL-MS (qXL-MS) can provide a structural understanding of protein dynamics (30–33). Here, we adopt XL-MS to study condensates.

In this study, we take a biochemical approach using purified proteins to reconstitute a chaperone-mediated quality control mechanism associated with RNP granules and combine it with qXL-MS to probe PPIs inside condensates. We find that unfolding of the RRM drives FUS aging and that interaction of its folded RRM with the small heat shock protein HspB8 slows down this aging process. Importantly, these condensate-specific interactions are altered in a disease-associated HspB8 mutant, resulting in its inability to prevent FUS aging.

## Results

### Quantitative and time-resolved XL-MS reveal domain-specific changes in crosslink abundances underlying condensate formation

To investigate the condensate specific interactions of FUS after phase separation, we diluted purified FUS-G156E protein (4), called FUS_m_ throughout this manuscript, either into a low salt solution, which induces phase separation or into a high salt buffer, which prevents phase separation. After crosslinking of lysine residues and subsequent digestion, equal amounts of peptides were subjected to mass spectrometry analysis to reveal condensate-specific crosslink patterns (**Figure 1A, Figure S1A, Supplementary Data 1**). These crosslinks can either reflect the spatial proximity of regions and protein-domains within a given protein, called **intra-links**, or between different proteins, called **inter-links**. For a detailed description of the different crosslink types *see* **Figure S1B**. When the crosslinker reacts twice within one peptide, this is called a **loop-link**. Additional information comes from cross linking of one side of the crosslinker with the protein and hydrolysis on the other side. This is called a **mono-link** and reveals information on the accessibility of a specific lysine residue. In some cases, we can also follow links between proteins of the same species (**homo-dimeric link**). These are crosslinks between overlapping peptides whose sequence is unique within the protein and that must therefore originate from different copies of the same protein.

**Figure 1.**
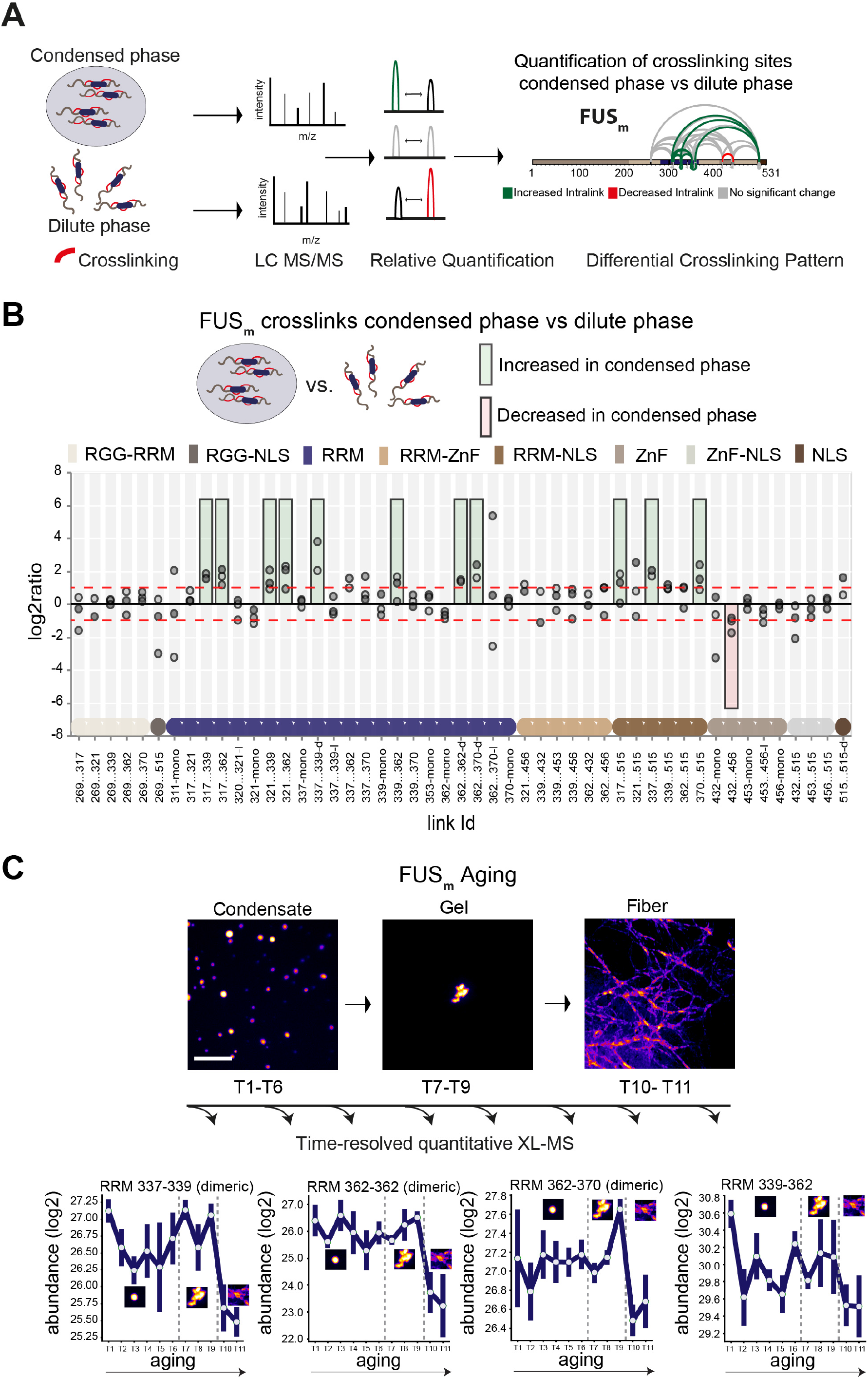
Domain-specific changes in crosslink abundances underlying condensate formation and molecular aging. (**A**) Workflow of quantitative crosslinking coupled to mass spectrometry (qXL-MS) of FUS_m_ condensates. (**B**) Crosslink abundance plot from reconstituted FUS_m_ condensates. Plotted are the relative enrichment (droplet vs non-droplet) for each unique crosslinking site (y-axis) sorted according to the known domain structure within FUS_m_ (x-axis). Shown are only high confidence crosslinking sites (see methods for details) from three biologically independent sets of experiments (n=3; circles in different shades of grey). Crosslinking sites that were consistently up – or downregulated two-fold or more (log2ratio ≥ 1 or ≤ −1 and FDR ≤ 0.05) in at least two out of three biological replicate sets and in addition contained no opposing regulation in any replicate set were considered significant and are highlighted with a green (enriched in droplets) or red background rectangle (decreased in droplets). All other changes in crosslinking abundances were considered insignificant and are shown on grey background. The significance threshold of two-fold enrichment is indicated as dashed red lines. Dimeric links are indicated by an additional “-d”, loop-links by a “-l” and mono-links by an “-mono” at their respective unique crosslinking site. (**C**) *Upper panel*: workflow time-resolved quantitative XL-MS. The conversion of fresh FUS_m_ condensates via the gel state into fibers was monitored by fluorescence microscopy. Scale bar is 10 µm. At indicated timepoints aliquots of the stock solution were crosslinked for 5 minutes, flash-frozen in liquid nitrogen and subsequently analyzed by MS (see methods for details). *Lower panel*: shown are changes of RRM crosslinks during aging that were increased during condensation (Figure 1B). The logarithmic total MS1 area for each time-point during aging is plotted (SDs; n=6). Domain structures within FUS_m_ are color-coded as in Figure 1B.

Please note that the crosslinker used throughout this manuscript, disuccinimidyl suberate (DSS), will react with primary amines and thus primarily link lysine residues within studied proteins. It is important to remember that the LCD of FUS contains no lysines and therefore our approach is not picking up interactions between and amongst the LCD domains.

The most prominent feature we detected after FUS_m_ condensation was that multiple intra-links within the RRM domain of FUS_m_ increased inside the condensates. This suggests that there are augmented contacts within the RRM domain, indicative of a structural change. In addition, there was an increase in homo-dimeric links between RRM domains, indicative of interactions between RRM domains of different FUS_m_ molecules. Additionally, links between the RRM domain and the nuclear localization sequence (NLS) were increased, while links within the Zinc-finger domain were decreased (**Figure 1B**). Taken together, these results suggest structural changes in the RRM domain accompany condensation.

### Crosslinks within the RRM domain change during molecular aging

It has been shown previously that FUS condensates undergo molecular aging in vitro, resulting in slowed down internal dynamics over time (4, 34, 35). To study protein dynamics and conformational changes during aging, we devised a time-resolved, quantitative XL-MS approach (**Figure 1C**) based on significantly shortened crosslinking times than conventionally used (36) (**Figure S1C**), which put us into the position to assess all stages of the aging process (**Figure S1D**). We then monitored the conversion of fresh FUS_m_ condensates into fibers by fluorescence microscopy (**Figure 1C**). At indicated timepoints the assemblies were crosslinked and analyzed by MS. We looked at 11 time points during the aging process and consistently quantified 77 crosslinks relative to fresh condensates (T1) (**Supplementary Data 2**). A global view of all quantified crosslinking sites shows that the vast majority of changes are happening during the formation of fiber states (**Figure S1E**). A closer examination focusing on those crosslinks within the RRM that were increased during condensation shows that these also change during molecular aging and particularly during fiber formation (**Figure 1C and S1F**). For instance, the link within the RRM bridging positions 339 and 362 decreases as fibers form. The general trend is for interactions between RRM domains and within RRM domains to decrease as condensates age.

### The small heat shock protein HspB8 partitions into FUS condensates and interacts with the RRM domain

To test whether these changes in links are indeed driven by aging, we next looked for ways to slow down the aging process. sHSPs are ATP-independent chaperones and are structurally divided into intrinsically disordered regions (IDRs) and a folded chaperone domain called α-crystallin domain (αCD) (37). HspB8 has previously been localized to stress granules (13). We therefore purified HspB8 (**Figure S2A**) and looked at its interaction with FUS_m_ condensates in vitro. We find that HspB8 was sequestered into reconstituted FUS_m_ condensates (**Figure 2A**) and that its fluorescence signal superimposed with the FUS_m_-GFP signal (**Figure 2B**), while it did not form droplets on its own under these conditions (**Figure S2B**). Another closely related small heat shock protein that has been localized to stress granules (12), HspB1, did not accumulate inside FUS_m_ droplets (**Figure 2C, Figure 2D, Figure S2A**). Using the fluorescence signal of labeled HspB8 and a calibration curve, we determined the concentration of HspB8 inside FUS_m_ condensates to be 2.4 mM (**Figures S2C & S2D**).

**Figure 2.**
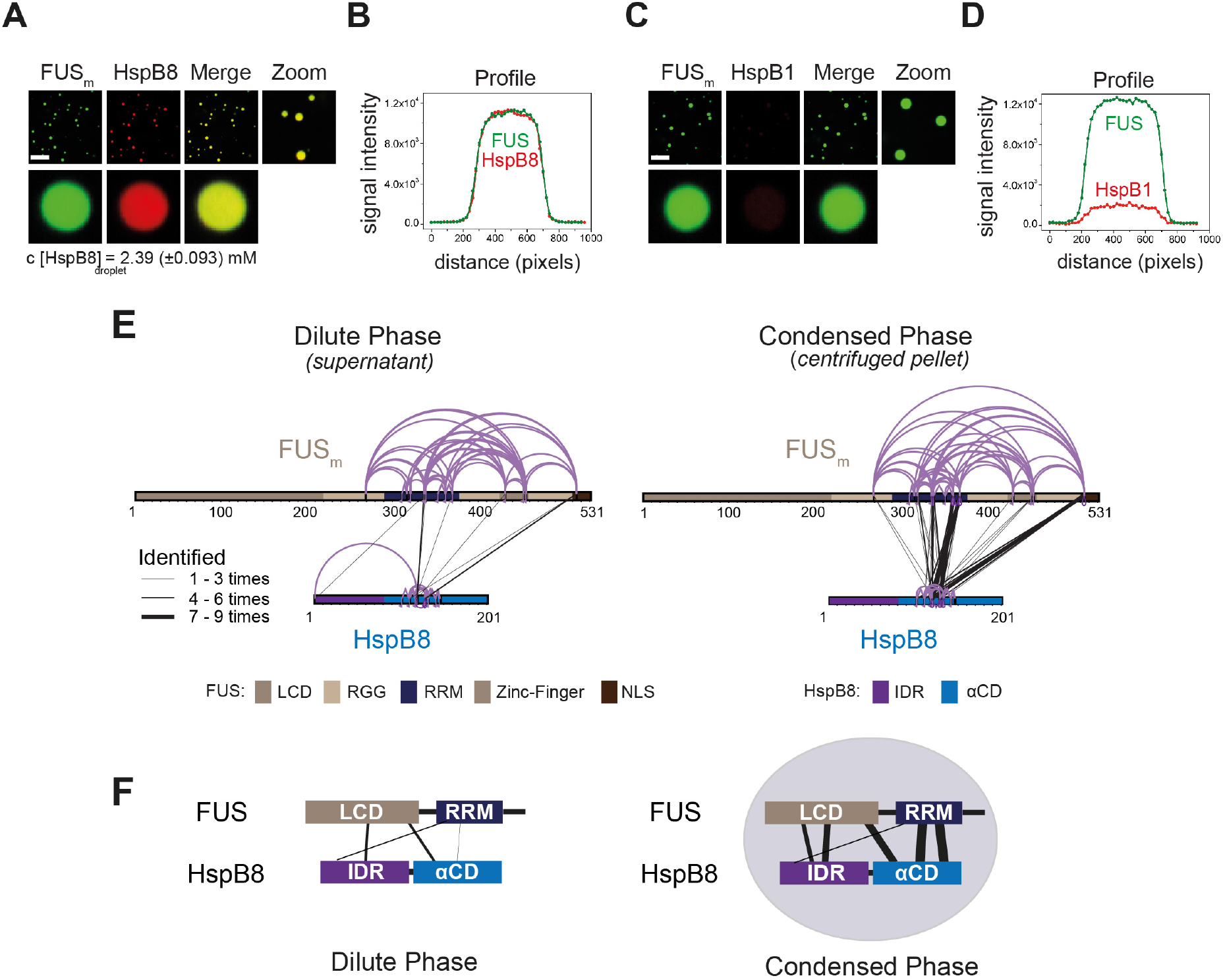
The small heat shock protein HspB8 partitions into FUS condensates and interacts with the RRM domain. (**A**) 0.25 µM Cy3-labelled HspB8, 4.75 µM unlabelled HspB8 (1:20 mix) and 5 µM FUS_m_-GFP were mixed in low salt buffer and the resulting droplets were imaged in a confocal microscope. Scale bar is 10 µm. (**B**) Fluorescence intensity plot profile spanning a FUS_m_-HspB8 droplet in (A). (**C**) 0.25 µM Cy3-labelled HspB1, 4.75 µM unlabelled HspB1 (1:20 mix) and 5 µM FUS_m_-GFP were mixed in low salt buffer and imaged in a confocal microscope. Scale bar is 10 µm. (**D**) Fluorescence intensity plot profile spanning a FUS_m_-HspB1 droplet in (C). (**E**) Overall crosslinking pattern of mixtures of FUS_m_ and HspB8 that were crosslinked under condensate-inducing low salt conditions (75 mM) and separated into the dilute phase (*left*) and condensed phase (*right*) by centrifugation. Experiments were carried out in three biologically independent sets of experiments (meaning separate batches of expressed protein). For one set of experiments each sample was independently crosslinked in triplicates and crosslinks were only considered, if they were identified in two out of three replicates with a deltaS < 0.95, a minimum Id score ≥ 20and an ld score ≥ 25 in at least one replicate (filtering was done on the level of the unique crosslinking site) and an FDR ≤ 0.05. Interlinks are shown in black and the total number of identifications is indicated by the thickness of the connection. Intralinks are shown in violet, monolinks with a flag, loop links with a pointed triangle and homodimeric links with a loop. (**F**) Representative overview of observed crosslinks between the LCD and RRM domains in FUS_m_ and the IDR and αCD domains in HspB8 in the dilute and the condensed phase as detected by XL-MS (based on Figure 2E, S2F and S2G).

We then used XL-MS, to probe protein-protein interactions (PPIs) between FUS_m_ and HspB8 inside the condensates. Inter-links (crosslinks between different proteins) predominantly formed inside the condensed phase (**Figure 2E, Supplementary Data 3**). Together with data from thermophoresis binding experiments (**Figure S2E**), this suggests that HspB8 specifically binds to FUS_m_ inside and not outside the condensed phase. We find the majority of inter-links within the droplets were formed between the αCD of HspB8 and the RRM of FUS_m_, and to a lesser degree the FUS-NLS. This condensate-specific and highly reproducible crosslinking pattern (**Figure 2E, Figure S2F & S2G**) was not seen when lactalbumin, a FUS-unrelated molten globule protein that unspecifically partitions into FUS_m_ condensates (**Figure S2H, S2I**) was used as a control. Taken together, interactions between HspB8 and FUS_m_ show a condensate-specific increase; in particular, interactions between the αCD of HspB8 and the FUS_m_-RRM are significantly upregulated inside condensates (**Figure 2F**).

### HspB8 prevents hardening and fiber formation of FUS droplets and keeps them dynamic

A time course experiment using photobleaching recovery (FRAP) revealed that in the presence of HspB8, FUS_m_ retained its liquidity over a time span of 24 hours (**Figure 3A to C, Figure S3A to D**). FUS_m_ droplets were able to fuse even after 6 hours (**Figure 3D**): the relaxation times of the fusion events were not affected (**Figure 3E**) and the drops no longer adhered to each other (**Figure S3E**). In addition to protecting fresh FUS_m_ condensates from converting into fibers (**Figure 3F**), the chaperone prevented further conversion of pre-aged FUS_m_ droplets, and prevented seeding of fiber growth (**Figure 3G, Suppl. Movie**). This correlated with the localization of HspB8 to FUS_m_ assemblies (**Figure S3F**). The protective effect of HspB8 on FUS_m_ was also observed at sub-stoichiometric HspB8 concentrations (**Figure 3H**).

**Figure 3.**
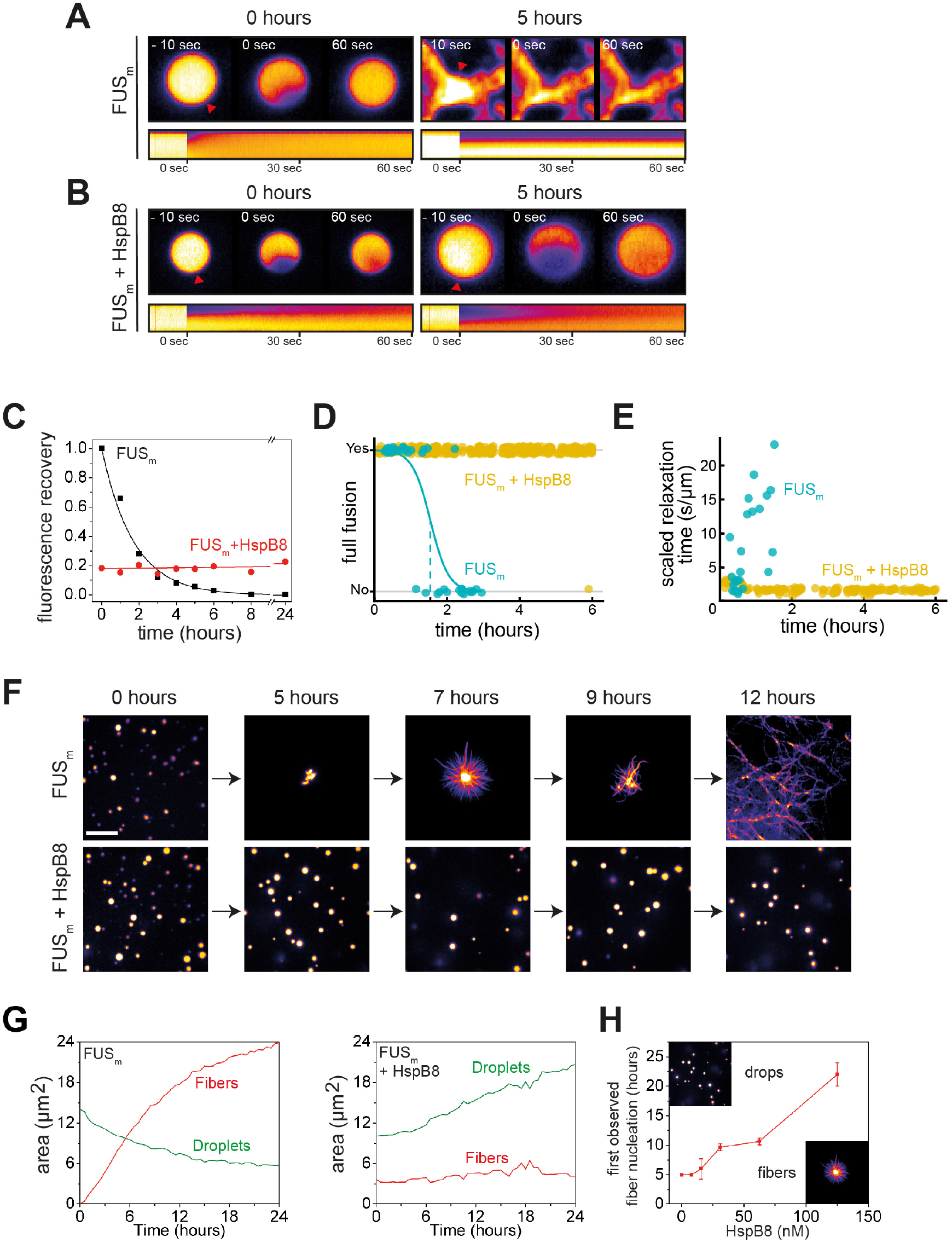
HspB8 prevents hardening and fiber formation of FUS droplets and keeps them dynamic. (**A**) FRAP experiment of fresh FUS_m_ condensates (0 hours) and condensates incubated for 5 hours. A kymograph shown below illustrates the kinetics of the process. (**B**) FRAP experiment of fresh FUS_m_ condensates mixed with HspB8 (0 hours) and condensates incubated for 5 hours. A kymograph shown below illustrates the kinetics of the process. (**C**) Kinetics of the FUS_m_ aging process. Plotted are the initial slopes of the FRAP recovery curves for FUS_m_ condensates in the absence (black) or presence of HspB8 (red). (**D**) Successful complete fusion events were registered over time, demonstrating the aging process of the FUS_m_ sample in the absence (turquoise, N = 40) or the presence of HspB8 (yellow, N = 330). The half-life of liquid-like FUS_m_ condensates alone was estimated to be around 1.5 h from logistic regression. (**E**) The size-normalized coalescence relaxation time is an indicator for the material state of the condensates. While it increases for FUS_m_ condensates during the hardening process, it stays constant over 6 hours in the presence of HspB8. Tweezer experiments were performed with fresh samples of 5 μM FUS_m_ with or without 20 μM HspB8. (**F**) Aging process of 5 µM FUS_m_ condensates in the absence and presence of 5 µM HspB8. In the presence of the chaperone the droplet morphology is maintained over the whole timeframe of the experiment (12 hours). Scale bar is 10 µm. (**G**) *Left panel:* shown is the total area of droplet material (green line) or fibrous material (red line) within FUS_m_ droplets as a function of time after FUS_m_ droplets were added to existing fibrous material. Spacing between data points is 30 min. *Right panel*: shown is the total area of droplet material (green line) and fibrous material (red line) within FUS_m_ droplets as a function of time after FUS_m_ droplets and HspB8 were added to existing fibrous material. Spacing between data points is 30 min. (**H**) The onset of 5 µM FUS_m_ fiber formation as a function of HspB8 concentration.

In order to investigate the molecular effect of HspB8 on the hardening of FUS_m_, we conducted quantitative XL-MS experiments in the presence of HspB8 and looked at crosslinks between FUS_m_ peptides. We found that previously observed crosslink patterns for pure FUS_m_ droplets, in particular upregulated crosslinks of the FUS-RRM, were still seen in the presence of HspB8 (**Figure S3G, Supplementary Data 4**). However, interactions between both the RRM with RGG and the RRM with the ZnF decreased, as did links within the ZnF and between the ZnF and the NLS. Our data show that HspB8 binds to the FUS-RRM also during prolonged incubation times via the αCD::RRM interface established during condensation (**Figure S3H, S3I**). This suggests that HspB8 binds to the RRM domain in condensates and prevents it from forming aberrant interactions with other domains in the protein.

### The disordered region of HspB8 directs the α-crystallin domain into FUS condensates for chaperoning

sHSPs consist of a αCD and flanking regions that are thought to be intrinsically disordered (38). Disorder prediction and circular dichroism analysis revealed that HspB8 is likely to be significantly more disordered than the closely related HspB1 (**Figure S4A-D**). To dissect the influence of the conserved αCD and the disordered IDR on the aging process of FUS_m_ we designed SNAP fusion constructs where we fused either only the IDR, the αCD or full length HspB8 to a fluorescently labelled SNAP-tag (**Figure 4A**). While the IDR-SNAP fusion construct was still recruited to FUS_m_ condensates, the αCD-SNAP variant no longer partitioned (**Figure 4B**). A decreased partitioning of the IDR-SNAP compared to the full-length construct indicated a contribution of the αCD (**Figure 4B, S4E**). The IDR-SNAP construct was also not active in preventing FUS_m_ fiber formation (**Figure 4C**), while the αCD-SNAP construct showed slight activity at high concentrations (**Figure S4F**). Similarly, in a FRAP assay the αCD-SNAP construct showed slight activity, while IDR-SNAP was inactive in preventing FUS_m_ gelation (**Figure S4G**). A partitioning analysis of these variants with droplets formed only by the LCD of FUS (AA1-211) under crowding conditions revealed a similar partitioning pattern as compared to full length FUS_m_ (**Figure S4H**), indicating that the HspB8-IDR interacts with the FUS-LCD. In order to rule out potential perturbations of HspB8 domain activity by fusion to the SNAP moiety, we designed additional swap variants of HspB8 and the closely related HspB1 (**Figure S4I**) and all our experiments with the HspB8-HspB1 swap variants mirrored the results with the HspB8-SNAP fusions (**Figure S4J–S4N**).

**Figure 4.**
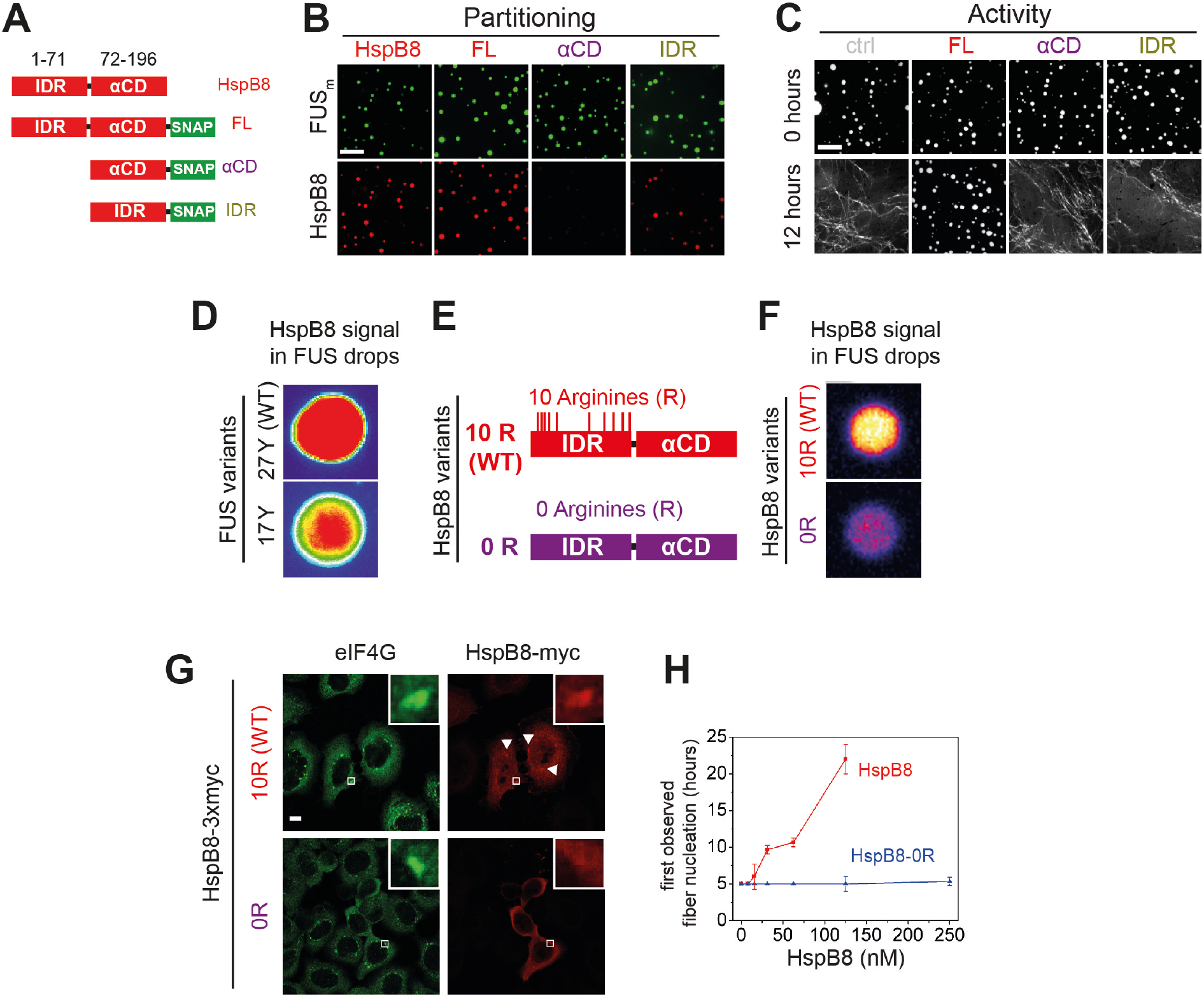
Arginines within the disordered region of HspB8 direct the α-crystallin domain into FUS condensates for chaperoning. (**A**) Overview of HspB8 truncation variants used in this study. FL (full-length HspB8 fused to a SNAP-tag), αCD (HspB8 alpha-crystallin domain (AA72-196) fused to a SNAP-tag), IDR (HspB8-N-terminus (AA1-71) fused to a SNAP-tag). (**B**) Partitioning of 5 µM HspB8 SNAP constructs (1:20 mix of labelled:unlabelled) into 5 µM FUS_m_ condensates. Scale bar is 10 µm. (**C**) Aging assay of 5 µM FUS_m_ condensates in the absence (ctrl) and presence of 5 µM HspB8 truncation variants. Scale bar is 10 µm. (**D**) Partitioning of 5 µM HspB8 (1:20 mix of labelled:unlabelled) into condensates formed by 5 µM FUS wildtype (27Y, WT) or a variant with a reduced number of tyrosines in its IDR (17Y). (**E**) Location of Arg residues in the primary structure of HspB8. 10 Arg residues are located in the N-terminus of HspB8-WT (10 R). In the HspB8-0R variant, these Arg residues are replaced by Gly residues (0 R). (**F**) Partitioning of 5 µM HspB8-WT or HspB8-0R variant (1:20 mix of labelled:unlabelled) into condensates formed by 5 µM FUS_m_. (**G**) HeLa Kyoto cells expressing HSPB8-WT-3xmyc or HSPB8-0R-3xmyc from a plasmid were subjected to heat shock at 43.5 °C for 1 hr. Cells were fixed and stained with myc and eIF4G specific antibodies. Merged image composed of eIF4G (green) and myc (red) signals is shown. (**H**) The onset of 5 µM FUS_m_ fiber formation as a function of HspB8-0R concentration.

In summary, these results suggest that the IDR of HspB8 targets the αCD to the condensed phase via interaction with the LCD of phase separated FUS and that the HspB8-αCD is the active domain in preventing FUS aging.

### Arginines in the disordered region of HspB8 direct the αCD into FUS condensates

Recent studies identified interactions between tyrosines in the LCD and arginines in the RBD of FUS to be crucial for its ability to phase separate (21, 22, 25). Because HspB8 partitions into condensates formed only by the FUS-LCD (**Figure S4N**), we hypothesized that HspB8 could interact with tyrosines in the LCD of FUS. We tested the partitioning of HspB8 into a variant of FUS with a decreased number of tyrosines in the FUS-LCD. FUS wildtype has 27 tyrosines in its LCD. Mutating all of these to serines in the 0 Y variant abrogates phase separation, but the 17 Y FUS variant still undergoes phase separation (22) (**Figure 4D**). Partitioning of HspB8 was significantly reduced for FUS condensates formed by the 17 Y variant (**Figure 4D**), suggesting that tyrosines in the FUS-LCD interact with HspB8. HspB8 has 10 arginines in its predicted IDR (**Figure 4E**). We exchanged these for glycines resulting in the 0 R variant of HspB8. This variant did not accumulate in FUS_m_ droplets in vitro (**Figure 4F**) and did not partition into stress granules in cells, contrary to the behavior of HspB8-WT (**Figure 4G**). When we tested the 0 R variant of HspB8 in a FUS_m_ aging assay we found that its ability to prevent fiber formation was completely abolished (**Figure 4H**).

### RRM unfolding drives FUS aging and is rescued by HspB8

Next, we sought to test consequences of the interaction between the HspB8-αCD and the FUS-RRM for FUS aging. To this end, we deleted the RRM in the variant FUS_m_ΔRRM (ΔAA285-371) and monitored its aging process in the presence and absence of HspB8 (**Figure 5A**). First, FUS_m_ aging was observed after 4 hours in the control condition, while co-incubation with HspB8 prevented the aging process over the time course of the experiment. Deletion of the RRM domain in FUS_m_ΔRRM significantly slowed down aging and first aggregates were observed after 36 hours. Remarkably, HspB8 was not able to prevent aging of the FUS_m_ΔRRM variant, indicating that HspB8 binding to the FUS-RRM is required for HspB8 to act as a chaperone for FUS. Our XL-MS data shows that the majority of inter-links between FUS_m_ and HspB8 that were significantly increased in the condensed phase in fact formed between the αCD of HspB8 and the RRM of FUS_m_ (**Figure 5B**). Thus, while the RRM domain does not seem to be solely responsible for FUS aging, it significantly contributes to the initiation of the process.

**Figure 5.**
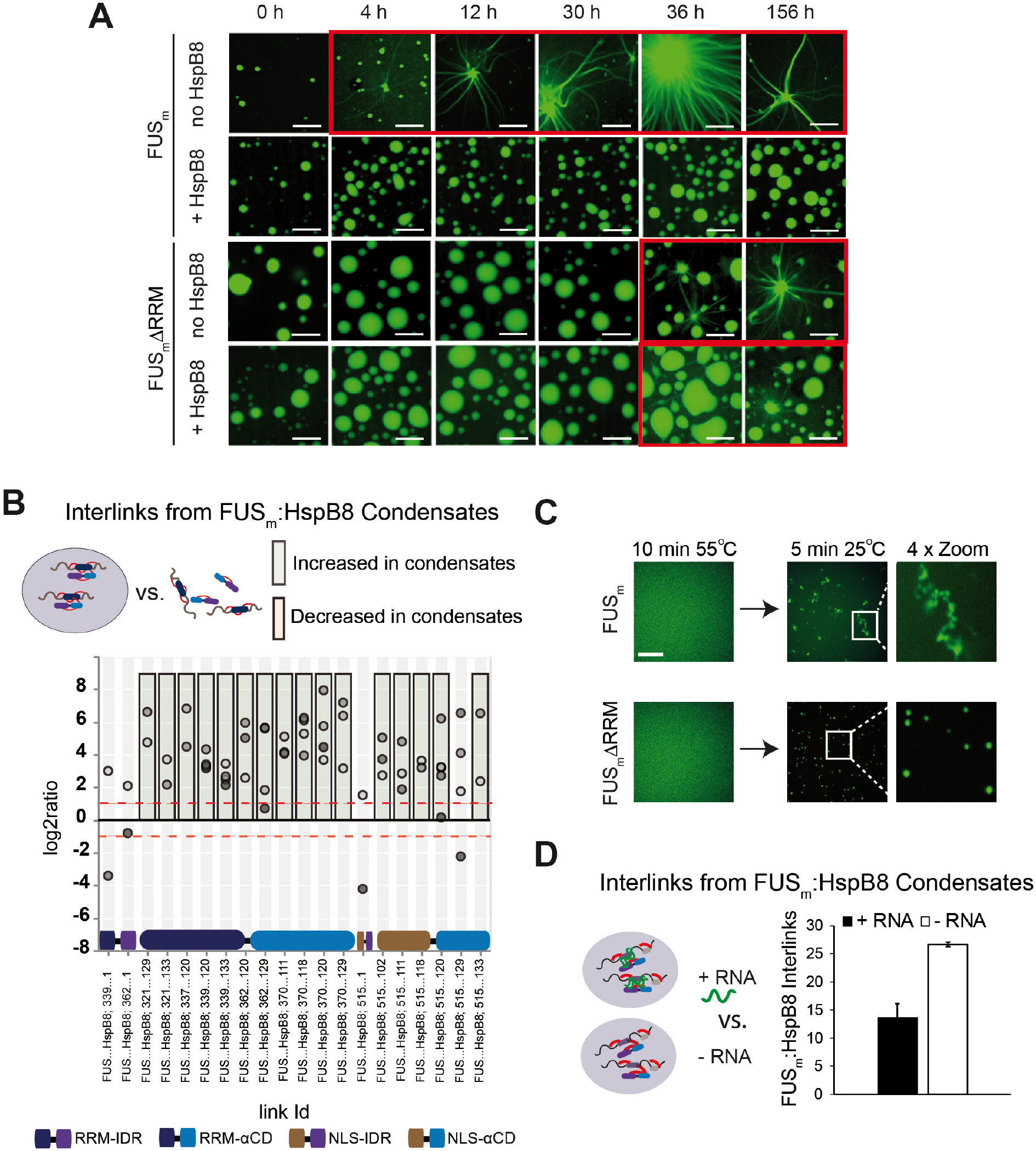
RRM unfolding drives FUS aggregation and is rescued by HspB8. (**A**) Aging process of molecular condensates containing either FUS_m_ or the FUS_m_ΔRRM (ΔAA285-371) variant in the presence or absence of HspB8 monitored by fluorescence microscopy over time. Scale bar is 10 µm. (**B**) Interlink abundances from reconstituted FUS_m_:HspB condensates. Plotted are the relative enrichment (droplet vs non-droplet) for each unique crosslinking site (y-axis) sorted according to the known domain structure within FUS_m_ and HspB8 (x-axis). Shown are only high confidence crosslinking sites (see methods for details) from five biologically independent sets of experiments (n=5; circles in different shades of grey). Crosslinking sites that were consistently up – or downregulated two-fold or more (log2ratio ≥ 1 or ≤ −1 and FDR ≤ 0.05) in at least two out of five biological replicate sets of experiments and in addition contained no opposing regulation in any replicate set were considered significant and are highlighted with a green (enriched in droplets) or red background rectangle (decreased in droplets). All other changes in crosslinking abundances were considered insignificant and are shown on grey background. The significance threshold of two-fold enrichment is indicated as dashed red lines. (**C**) Fluorescence microscopy images of FUS_m_ and FUS_m_ΔRRM variant during and after incubation under heat-shock conditions. Samples were heated to 55°C for 10 min followed by a 5 min cool-down step to 25°C. Scale bar is 10 µm. (**D**) FUS_m_:HspB8 condensates were crosslinked in the presence (black) or absence (white) of a customized RNA oligonucleotide previously shown to bind to FUS (40) and analysed by LC MS/MS (n=3; FDR ≤ 0.05).

The RRM domain of FUS is a folded domain in an otherwise disordered protein (**Figure S5A**). We suspected that RRM unfolding might serve as a seed for the formation of FUS aberrant conformations that would initiate FUS aggregation and fiber formation. To test this hypothesis, we performed a heat shock experiment to unfold the RRM domain of FUS (**Figure 5C**). The melting temperature of the isolated FUS-RRM has been reported to be 52 °C (29). We prepared reactions of condensates formed by full-length FUS or FUS_m_ΔRRM and incubated these for 10 min at 55°C to unfold the RRM. At this temperature FUS_m_ condensates were dissolved (**Figure 5C**). We then cooled down the reactions to 25°C and assessed the reactions by fluorescence microscopy. While full-length FUS_m_ formed amorphous aggregates after cooling down from heat shock, the FUS_m_ΔRRM variant condensed into spherical droplets (**Figure 5C**). This result strongly suggests that unfolding of the RRM domain is an integral part of the aging process and deletion of the RRM prevents temperature induced aggregation of FUS_m_ condensates.

It has been shown that RNA can bind to the FUS-RRM, prevent aging of FUS and dissolve condensates at high concentrations (39, 40). Hence, we were wondering whether RNA exerts these effects by virtue of binding and stabilizing the RRM domain. We tested this by looking at competition between RNA and HspB8 binding to FUS_m_ using XL-MS and crosslinked condensates of FUS_m_ and HspB8 in the presence or absence of RNA. While the addition of RNA led to a significant decrease in the number of detected interlinks between FUS_m_ and HspB8, it did not alter the number of intra-links within FUS_m_ or HspB8 (**Figure 5D & S5B, Supplementary Data 5**). This result strongly suggests that RNA and HspB8 compete for binding to FUS_m_ and indicate a similar mechanism by which they stabilize its RRM domain to prevent FUS_m_ aging.

### A disease-associated mutation interferes with HspB8 activity

Mutations of the lysine 141 residue in the αCD of HspB8 have been associated with Charcot-Marie Tooth disease (CMT), a currently incurable dominant autosomal disorder of the peripheral nervous system leading to muscular dystrophies (41) (**Figure 6A**). The mechanistic cause of the disease is still enigmatic, although experimental evidence indicates a decreased chaperone activity for HspB8-K141E (42–44). We introduced the K141E mutation into HspB8 and when we tested HspB8-K141E for localization to FUS_m_ droplets, we found that it still partitioned into reconstituted FUS_m_ droplets (**Figure 6B**). While the HspB8 wildtype (WT) was active in a FUS_m_ aging experiment, HspB8-K141E could not prevent FUS_m_ aging (**Figure 6C**) and when mixed with the WT, the mutant exerted a dominant negative effect over the WT, preventing the WT from being active. By using FRAP, we found that the mutant was much less effective compared to the WT in keeping FUS_m_ in a dynamic state (**Figure 6D**). Remarkably, the WT-mutant mix showed even lower chaperone activity, underlining the dominant negative role of HspB8-K141E mutant over the WT (**Figure 6D**).

**Figure 6.**
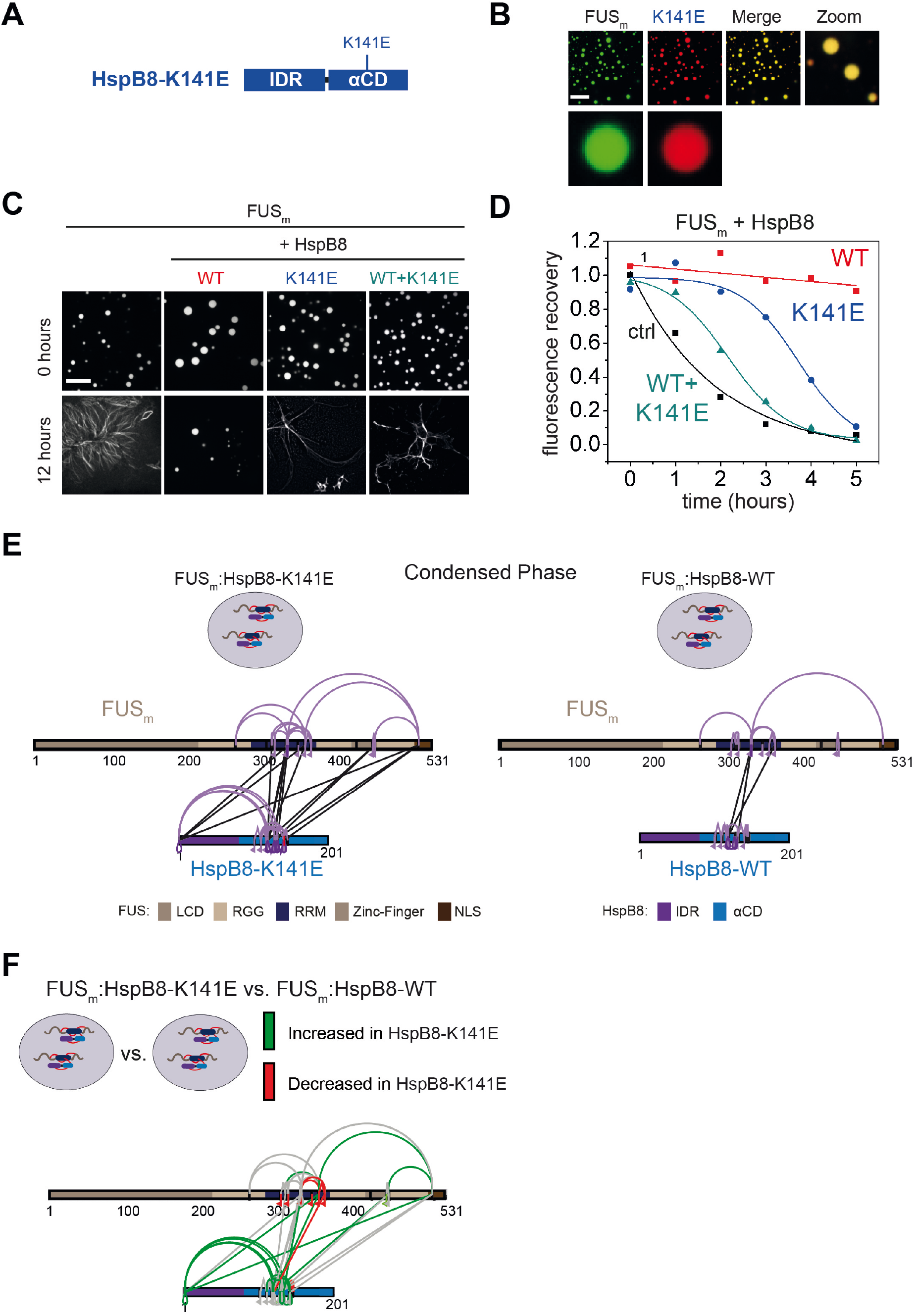
A disease-related mutation interferes with HspB8 activity. (**A**) Location of the disease related K141E mutation inside the αCD of HspB8. (**B**) Partitioning of 5 µM HspB8-K141E (1:20 mix of labelled:unlabelled) into condensates formed by 5 µM FUS_m_. Scale bar is 10 µm. (**C**) Aging assay of 5 µM FUS_m_ condensates in the absence (ctrl) and presence of HspB8-WT, HspB8-K141E or a 1:1 mix of WT and K141 mutant. Final chaperone concentrations in all reactions are 125 nM. Scale bar is 10 µm. (**D**) Kinetics of the aging process. Plotted are the initial slopes of the FRAP recovery curves for FUS_m_ condensates in the absence (ctrl) and presence of HspB8-WT (red), HspB8-K141E (blue) or a 1:1 mix of WT and K141E mutant (cyan). (**E**) Equal amounts of FUS_m_:HspB8-K141E (left) or FUS_m_:HspB8-WT (right) were crosslinked under condensate inducing low salt conditions (75 mM NaCl). Experiments were carried out in triplicates and crosslinks were only considered, if they were identified in two out of three replicates with a deltaS < 0.95, a minimum Id score ≥ 20 and an ld score ≥ 25 in at least one replicate (filtering was done on the level of the unique crosslinking site) and an FDR < 0.05. The mutated site in HspB8-K141E is shown in red. Interlinks are shown in black and intralinks are shown in violet. Monolinks are shown with a flag, loop links with a pointed triangle and homodimeric links with a loop. (**F**) Quantitative comparison of crosslinking pattern from HspB8-K141E and HspB8-WT condensates. Crosslinking sites that were up – or downregulated two-fold or more (log2ratio ≥ 1 or ≤ −1 and FDR ≤ 0.05) were considered significant and are highlighted in green (i.e. relative enrichment in FUS_m_:HspB8-K141E condensates) or red (i.e. relative decrease in FUS_m_:HspB8-K141E condensates). All other changes in crosslinking abundances were considered insignificant and are shown in grey background.

We then performed XL-MS using the HspB8-K141E mutant. In comparison to HspB8-WT, the mutant showed an increased number of inter-links with FUS_m_ in the condensed phase, which suggests a stronger interaction between FUS_m_ and the HspB8-K141E mutant (**Figure 6E, Supplementary Data 6**). A quantitative analysis revealed that almost all mono-links within the RRM domain of FUS_m_ were decreased upon binding of the HspB8-K141E mutant, suggesting that accessibility to these sites was hindered (**Figure 6F, Supplementary Data 6**). Concomitantly, multiple intra-links within HspB8 bridging the N-terminus to the αCD domain were increased in the HspB8-K141E mutant (**Figure 6F**), an observation that is in line with a potential conformational change of the chaperone mutant upon FUS binding that brings these domains closer to each other (31). We conclude that the CMT patient mutations in a critical residue of HspB8 may lock interactions with FUS, thereby interfering with the dynamic interplay between the chaperone mutant and its client, which in turn may impair its ability to maintain FUS in a dynamic state.

## Discussion

In this study we show that the molecular chaperone HspB8 can prevent a disease-associated aberrant phase transition that is mediated by the protein FUS. We show that HspB8 uses its disordered domain to partition into liquid FUS condensates and that HspB8 uses a similar molecular grammar as described previously for FUS (22). More specifically, arginine residues in the IDR of HspB8 interact with tyrosine residues in the FUS-LCD, thereby promoting the targeting of the HspB8-αCD into FUS condensates for chaperoning of the misfolding-prone RRM domain of FUS. This suggests a general principle for how the protein quality control machinery could be targeted to condensates in order to regulate misfolding-prone protein domains inside liquid condensates.

Despite the extensive work both in vivo and in vitro on proteins that phase separate, our current knowledge on how proteins are organized within condensates is limited and monitoring the transient interactions inside condensates has remained a major challenge to the field. In principle, proteomics and mass spectrometry should be an appropriate method to map condensate specific interactions but reports using MS to study condensates remain scarce (28, 45, 46). Recent studies demonstrate that proximity labelling in combination with MS is well suited to track the protein content of a specific molecular condensate (10, 11), but falls short in charting direct protein-protein interactions or determining exact interaction sites. We and others showed previously that XL-MS is well suited to map PPIs and that relative changes in crosslinking as probed by quantitative XL-MS can provide a structural understanding of protein dynamics (30–33, 47). Here, we adopt XL-MS to study condensates. In doing so we show with unprecedented molecular detail how protein contacts are formed within molecular condensates and demonstrate for the first-time condensate-specific client interaction.

Although the role of small heat shock proteins in maintaining correct protein folding is intensively studied, up to now, the substrates of HspB8 have remained enigmatic (48). Our crosslinking data suggests that HspB8 exerts its effect in part through the FUS-RRM domain. Misfolding of the RRM domain may represent the initial step on the pathway to forming a seed that subsequently promotes the nucleation of FUS fibrils. This may involve cross-beta sheet interactions of the LCDs via local concentration and LCD alignment. In this model, HspB8 would stabilize the fold of the RRM domain by binding to it, and by doing so maintain the liquid state of the condensates (**Figure 7**).

**Figure 7.**
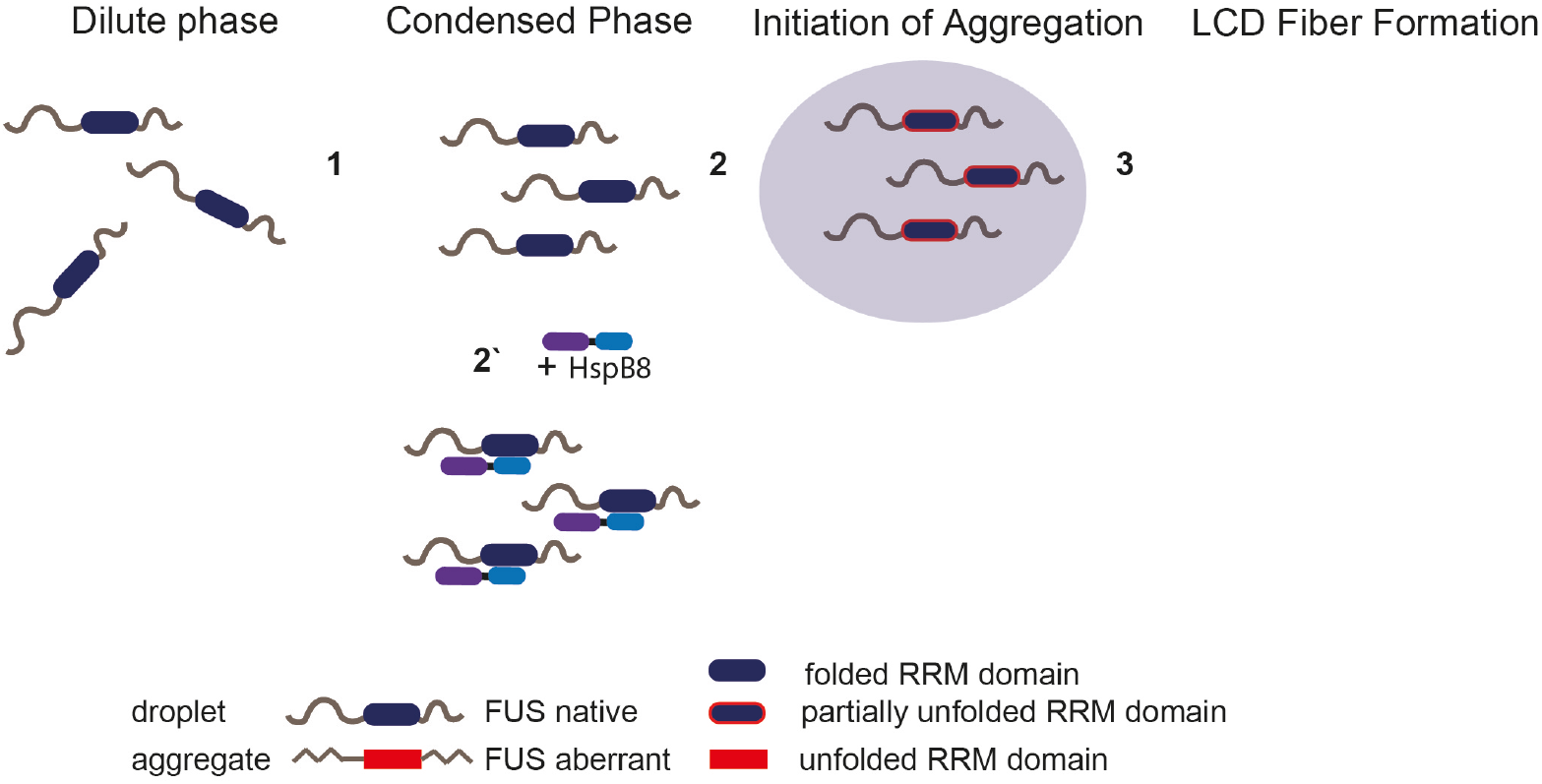
Unfolding of globular domains nucleates aberrant seeds for aggregation. Proposed mechanism where unfolding of globular domains in otherwise disordered proteins nucleates aberrant seeds for aggregation. In this model the unfolding of the RRM domain comprises the initial step to seed the conversion of FUS into an aberrant conformational space that consequently involves the beta-amyloid formation of the LCDs by rising their local concentration and potentially by sterically aligning them for nucleation. Molecular chaperones like HspB8 stabilize the fold of the RRM domain and maintain the dynamic liquid state of the condensate.

RRM domains are small folded entities that bind to single-stranded RNA. In human, there are currently 745 RRM domains distributed over 446 proteins known to exist (49). It has been proposed before that cells use RNA to regulate condensate formation (40). We hypothesize that some RRM domains require stabilization by binding to RNA and in cases where no RNA is available, the chaperone HspB8 takes over this function and protects the RRM domain from unfolding and aggregation. Thus, it is a possibility that HspB8 has a general role in cell physiology in stabilizing RRM domains in condensates.

The changes we find in the inter-link patterns between FUS_m_ and HspB8-WT or the neuropathy-causing mutant HspB8-K141E point to shifts in conformation and dynamics of the RRM domain after condensation. These shifts may differentially affect the fate of the bound substrate: stabilization of the FUS_m_ native conformation by HspB8-WT versus loss of function by the HspB8-K141E mutant with subsequent FUS_m_ aggregation. To date, mutations in the genes coding for HspB1, HspB3, HspB5 and HspB8 have been associated to neuromuscular diseases (18). To what extent similar mechanisms may also affect other HspB mutants therefore awaits further investigation.

A mechanism by which unfolding of globular domains is controlled by chaperones inside condensates may be a process of general importance (50). Deletion of the RRM slows down hardening but does not stop it. This suggests other driving forces such as aberrant LCD-LCD interactions that may require other types of chaperone systems to prevent them. Our study also highlights the interplay between folded domains and IDRs. A changed environment after phase separation could destabilize folded domains and require chaperones to stabilize them. Such a mechanism would necessitate many domain-specific chaperones for stabilization inside condensates.

In summary, by adapting existing XL-MS workflows, we were able to monitor PPIs and protein dynamics inside reconstituted protein condensates, thus paving the way for a deeper and more detailed structural understanding of condensate formation and aberrant phase transitions. Our study shows with unprecedented molecular detail how protein contacts are formed between a chaperone and the folded RRM of its client protein inside a condensate and therefore suggest a blueprint for how chaperones could act to stabilize biomolecular condensates in cells. More generally, our data suggests that established principles of cellular organization, such as domain-specific protein-protein interaction sites, also apply to the biochemistry inside molecular condensates. Here, further work is required to expand the resolution of crosslinking mass spectrometry by developing crosslinking chemistries optimized for intrinsically disordered domains and to augment its current ability to also target condensates in vivo.

## Supplementary Figures

**Figure S1.**
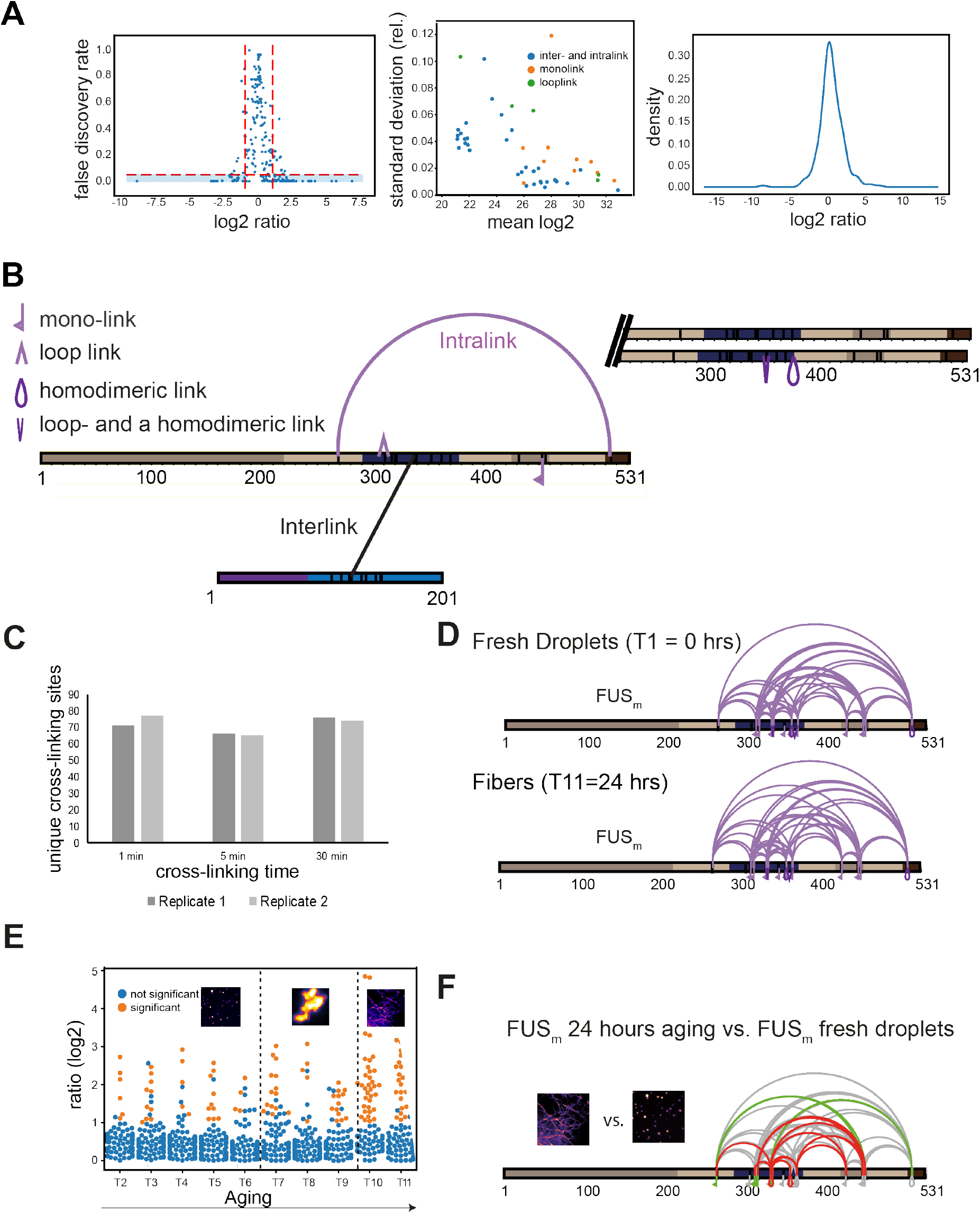
Related to Figure 1. (**A**) Statistical validation of crosslinking data. *Left panel* shows the relationship between the log2ratio and FDR for qXL-MS data. FDR values are p-values corrected for multiple testing (see methods for details). Significant crosslinks are shown in the blue-shaded area (log2ratio ≥ 1 or ≤ −1 and FDR ≤ 0.05). The majority of crosslinks with a high FDR value are found between the log2ratios of −1 and +1. The *middle panel* shows that quantifications are highly reproducible within triplicate measurements. Plotted is the absolute mean MS1 peak area per crosslink and experiment in log2-scale (where the mean is made up of the light and heavy state and the biological replicates of a crosslink) (x-axis) versus the relative standard deviation of the mean (y-axis). It increases with a decreasing mean but remains below 10% for virtually all crosslinks. The *right panel* shows that the overall distribution of log2ratios is centered around 0, confirming that the data has no systematic bias. (**B**) Disuccinimidyl suberate (DSS) was used to introduce covalent bonds between proximal primary amines in order to link lysine residues within studied proteins. The actual crosslinking sites are subsequently identified by MS and reflect the spatial proximity of regions and protein-domains within a given protein (**intra-link**) or between different proteins (**inter-link**). Additionally, the crosslinker can react twice within one peptide (**loop-link**) or only on one side with the protein and hydrolyze on the other side (**mono-links**), revealing information on the accessibility of a specific lysine residue. XL-MS cannot readily discriminate if a crosslink has formed within one polypeptide chain (defined as **intra-link**, vide supra) or between homodimers or even higher oligomers of the same protein. This is usually not a problem, but in the case of the high protein concentrations within the condensed phase it may play a role. However, as we detect all shown intra-links within FUS_m_ and HspB8 also under the relatively low-concentration regime of the dilute phase, it is fair to assume that all of the links are indeed intra-links as usually defined - e.g. occurring within one FUS_m_ polypeptide chain - as we assume in the current version of the manuscript. For some specific intra-links, we do however know that they must have occurred between different molecules of the same protein; these are crosslinks between overlapping peptides whose sequence is unique within the protein and that must therefore originate from different copies of the same protein (**homodimeric link**). (**C**) Significantly shorter crosslinking time periods than conventionally used lead to reproducible and sufficient crosslinking yields. A test protein was crosslinked for different time periods (1 min, 5 min and the standard 30 min) before the reaction was quenched and crosslinks were identified by LC MS/MS (data for two biological replicates are shown). In order to minimize the induced error rate due to sample handling, a crosslinking period of 5 minutes was chosen for all timepoints. (**D**) Overall crosslinking pattern of FUS_m_ from fresh droplets (T1 = 0 hrs) (upper panel) and the final fiber state (T11 = 24 hrs) (lower panel) reveal a similar overall crosslinking pattern. Experiments were carried out in triplicates and crosslinks were only considered, if they were identified in two out of three replicates with a deltaS < 0.95, a minimum Id score ≥ 20 and an ld score ≥ 25 in at least one replicate (filtering was done on the level of the unique crosslinking site) and an FDR < 0.05. (**E**) Overview plot of all crosslinks consistently quantified over 11 timepoints during the aging process. Crosslinks with a significant change relative to fresh condensates (T1) (log2ratio ≥ 1 or ≤ −1 and FDR ≤ 0.05) were colored orange while non-significantly changed crosslinks were colored blue. (**F**) Comparison of the overall differential crosslinking pattern from FUS_m_ from the final fiber state (T11 = 24 hrs) versus fresh droplets (T1 = 0 hrs). Crosslinking sites that were up – or downregulated two-fold or more (log2ratio ≥ ±1.0 and FDR ≤ 0.05) were considered significant and are highlighted in green (i.e. relative enrichment in final fiber state) or red (i.e. relative decrease in final fiber state). All other changes in crosslinking abundances were considered insignificant and are shown in grey background.

**Figure S2.**
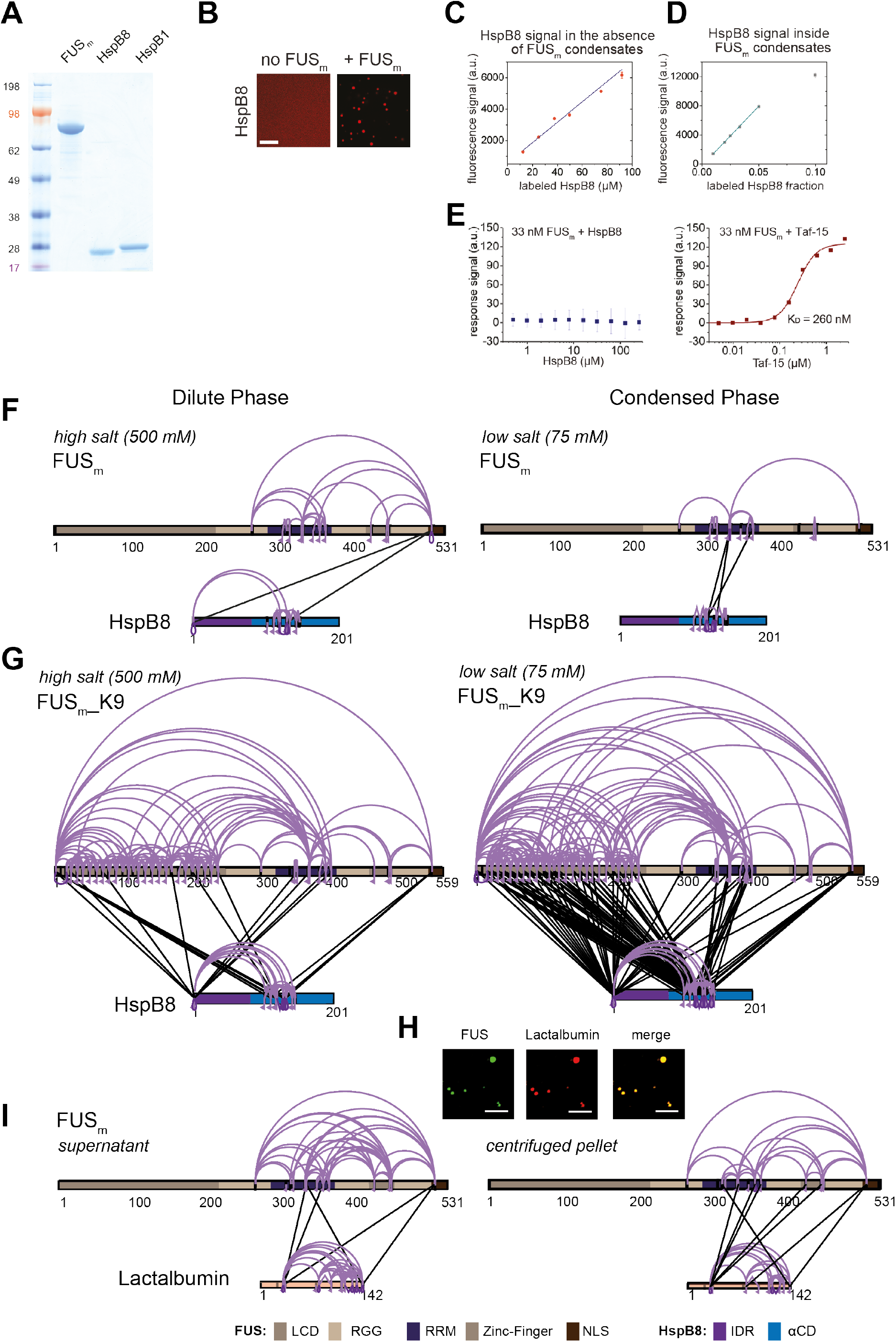
Related to Figure 2. (**A**) SDS-PAGE of purified FUS_m_, HspB8 and HspB1 proteins used in this study. (**B**) 5 µM HspB8 (1:20 mix of Cy3-labelled:unlabelled) in the absence and presence of 5 µM FUS_m_, Cy3 channel is shown. Scale bar is 10 µm. (**C**) Calibration curve for pure Cy3-labelled HspB8 in the absence of condensates. (**D**) Cy3 fluorescence signal inside condensates formed by 5 µM FUS_m_ in the presence of different ratios of Cy3-labelled and unlabelled HspB8. The Cy3 fluorescence signal inside the condensates is linear up to a ratio of 1:20 of labelled:unlabelled HspB8 (labelled HspB8 fraction of 0.05). (**E**) To test for a potential interaction between HspB8 and FUS_m_ in the dilute phase, protein-protein interactions of 33 nM FUS_m_ and HspB8 (*left panel*) and Taf-15 (*right panel*) were assessed under non-phase separating conditions (500 mM KCl) by Microscale Thermophoresis (51). While FUS_m_ tightly bound to the stress granule protein Taf-15 with a K_D_ of 260 nM, there was no detectable interaction between HspB8 and FUS_m_, even at a chaperone concentration of 120 µM. (**F**) Overall crosslinking pattern from mixtures of recombinant FUS_m_ and HspB8 under identical high salt (500 mM NaCl) (*left*) and condensate inducing low salt (75 mM NaCl) (*right*) conditions as used for the fluorescence microscopy experiments (Figure 2A and 2C). (**G**) Overall crosslinking pattern of mixtures of the recombinant FUS_m__K9 variant and HspB8 under high salt (500 mM KCl) (*left*) and condensate inducing low salt (75 mM KCl) (*right*) conditions. As FUS_m_ contains no lysine residues within its LCD which are amenable to our NHS-ester crosslinker, we generated a FUS_m_ variant with an additional “26” lysines in the LCD (FUS_m__K9), which resembles FUS_m_ in its condensation behavior even though it does not exhibit a similar ageing phenotype. We find that in the condensed phase the number of inter-links between the LCD of FUS_m_ and both the IDR and the αCD of HspB8 was strongly increased, confirming previous results on the role of the LCD during phase separation (22). However, only inter-links between the αCD of HspB8 and the RRM of FUS_m_ were exclusively detected inside the condensed phase. (**H**) Microscopic images of FUS_m_ and lactalbumin under condensate inducing high salt conditions (500 mM NaCl). Scale bar is 10 µm. (**I**) As a control we investigated the interaction between FUS_m_ and lactalbumin. Lactalbumin is a molten globule protein that partitions into FUS_m_ condensates likely mediated by unspecific interactions between partially exposed hydrophobic patches in the client and the condensate scaffold protein. Mixtures of FUS_m_ and lactalbumin that were crosslinked under condensate inducing low salt conditions (75 mM) and that were separated into the dilute phase (*left*) and condensed phase (*right*) by centrifugation did not show any significant differences in the overall crosslink patterns of dilute and condensed phase. This suggests that in contrast to the HspB8: FUS_m_ interaction, partitioning of lactalbumin into FUS_m_ condensates is caused by unspecific interactions. (**F, G and I**) Samples were independently crosslinked in triplicates and crosslinks were only considered, if they were identified in two out of three replicates with a deltaS < 0.95, a minimum Id score ≥ 20 and an ld score ≥ 25 in at least one replicate (filtering was done on the level of the unique crosslinking site) and an FDR ≤ 0.05. Interlinks are shown in black, Intralinks are shown in violet, monolinks with a flag, loop links with a pointed triangle and homodimeric links with a loop. Cases where a looplink and a homodimeric link were identified on the same lysine are indicated by a triangle pointed downwards. LCD, RGG, RRM, ZnF and NLS of FUS_m_ are shown in different shades of brown and blue, respectively. The IDR and αCD in HspB8 are coloured in dark violet and dark blue, respectively.

**Figure S3.**
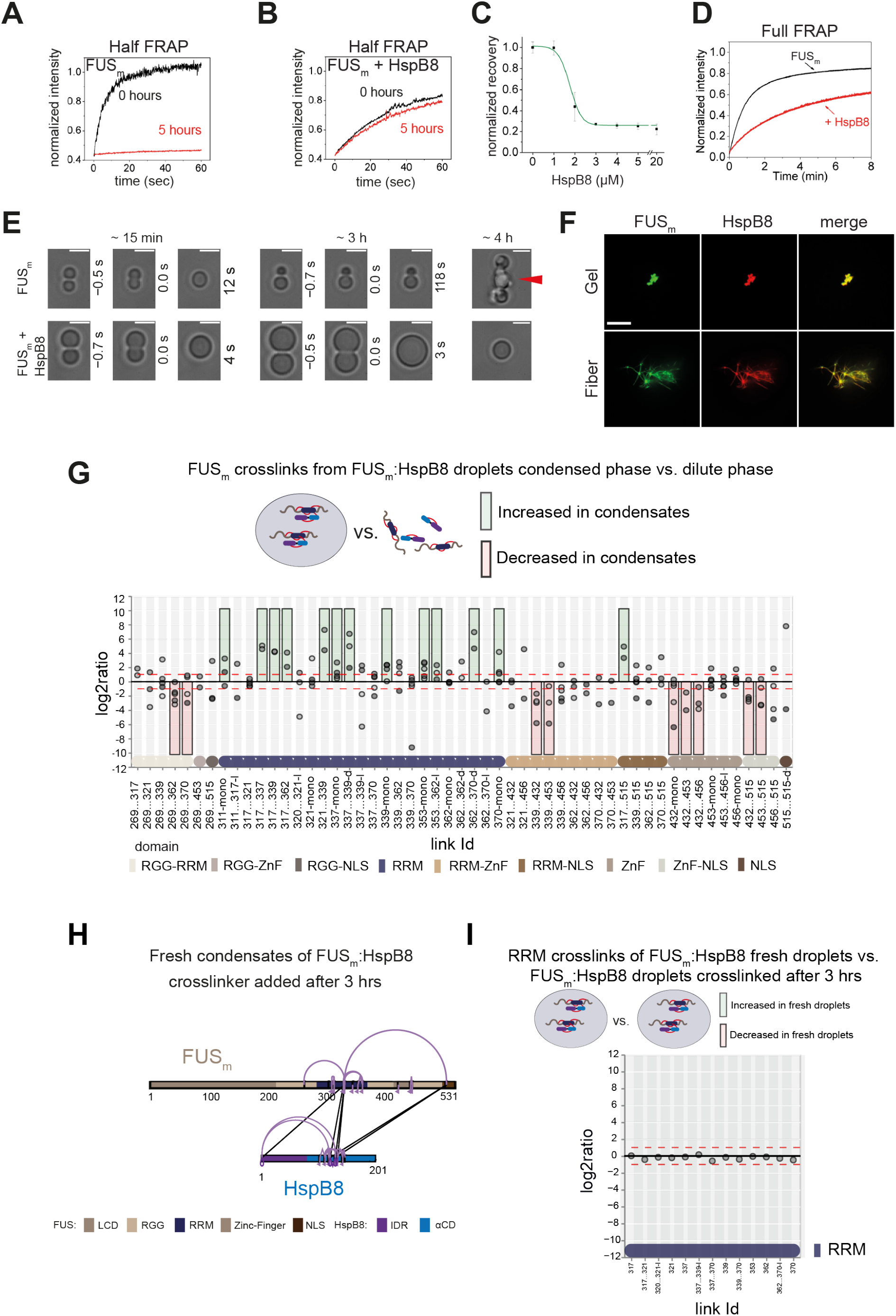
Related to Figure 3. (**A**) Quantification of a half-bleach FRAP experiment of fresh FUS_m_ condensates (black) and condensates incubated for 5 hours (red). (**B**) Same experiment as in (A) in the presence of 5 µM HspB8. (**C**) HspB8-dependent decrease in mobility has an EC50 of 1.5 µM. Plotted are the initial slopes of the FRAP recovery curves for 5 µM FUS_m_ condensates in the presence of different concentrations of HspB8. (**D**) Quantification of a full-bleach FRAP experiment of fresh FUS_m_ condensates in the absence (black) and presence of 5 µM HspB8 (red). (**E**) Representative images of attempted fusion events at early and late stages. Attempts to fuse several droplets of FUS_m_ condensates after hardening result in large amorphous assemblies (red arrow). Scale bar: 3 μm. (**F**) Colocalization of gels or fibers formed by FUS_m_ and Cy3-labeled HspB8. Scale bar is 10 µm. (**G**) Crosslink abundance plot from reconstituted FUS_m_:HspB8 condensates. Plotted are the relative enrichment (droplet vs non-droplet) for each unique crosslinking site (y-axis) sorted according to the known domain structure within FUS_m_ and HspB8 (x-axis). Shown are only high confidence crosslinking sites (see methods for details) from five biologically independent sets of experiments (n=5; circles in different shades of grey). Crosslinking sites that were consistently up – or downregulated two-fold or more (log2ratio ≥ 1 or ≤ −1 and FDR ≤ 0.05) in at least two out of five biological replicate sets of experiments and in addition contained no opposing regulation in any replicate set were considered significant and are highlighted with a green (enriched in droplets) or red background rectangle (decreased in droplets). All other changes in crosslinking abundances were considered insignificant and are shown on grey background. The significance threshold of two-fold enrichment is indicated as dashed red lines. Dimeric links are indicated by an additional “-d”, loop-links by a “-l” and mono-links by an “-mono” at their respective unique crosslinking site. (**H**) Overall crosslinking pattern of FUS_m_:HspB8 condensates that were allowed to mature and that were crosslinked 3 hours after their initial formation. (**I**) Focus on the RRM domain. A comparison of the crosslinks within the RRM (intra, loop and mono-links) shows no change in abundance when FUS_m_:HspB8 condensates of fresh droplets were compared with those that were crosslinked 3 hours after their initial formation.

**Figure S4.**
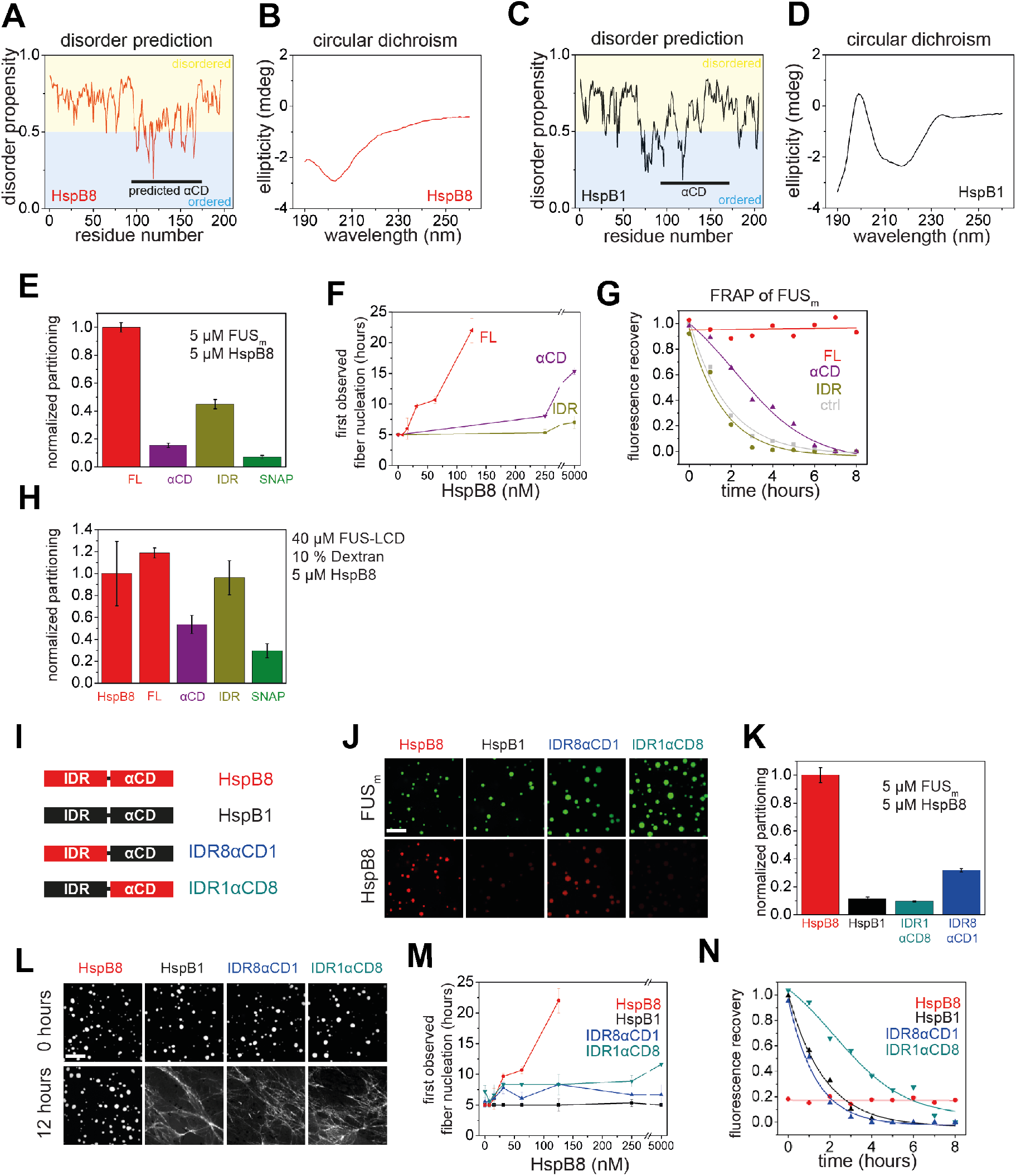
Related to Figure 4. (**A to D**) Disorder prediction and circular dichroism of HspB8 and closely related HspB1. (**A**) Disorder prediction plot using Dis-EMBL1.5 (52) for HspB8 shows that except for the αCD a large portion of the chaperone is predicted to be disordered while (**C**) HspB1 is predicted to be much less disordered. The αCD of HspB8 was predicted by sequence similarity to the crystallized αCD of HspB1. (**B**) Circular Dichroism analysis shows the typical spectrum of a disordered protein for HspB8, while the analysis of HspB1 (**D**) reveals mainly beta-sheet content. (**E**) Quantification of the partitioning of HspB8 truncation variants into condensates formed by full-length FUS_m_ as shown in (Fig. 4B). (**F**) The onset of 5 µM FUS_m_ fiber formation as a function of HspB8 truncation variant concentration. (**G**) Kinetics of the FUS_m_ aging process. Plotted are the initial slopes of the FRAP recovery curves for FUS_m_ condensates in the absence and presence of 5 µM HspB8 truncation variants. (**H**) Quantification of the partitioning of HspB8 truncation variants into condensates formed by the FUS-LCD in the presence of 10 % Dextran. (**I**) Overview of HspB8-HspB1 swap variants used in this study. IDR8αCD1 carries an IDR from HspB8 and an αCD from HspB1. IDR1αCD8 carries an IDR from HspB1 and an αCD from HspB8. (**J**) Partitioning of 5 µM HspB8-HspB1 swap variants (1:20 mix of labelled:unlabelled) into 5 µM FUS_m_ condensates. Scale bar is 10 µm. (**K**) Quantification of the partitioning experiment described in (G). (**L**) Aging assay of 5 µM FUS_m_ condensates in the absence (ctrl) and presence of 5 µM HspB8 truncation variants. Scale bar is 10 µm. (**M**) The onset of 5 µM FUS_m_ fiber formation as a function of HspB8-HspB1 swap variant concentration. (**N**) Kinetics of the FUS_m_ aging process. Plotted are the initial slopes of the FRAP recovery curves for FUS_m_ condensates in the absence and presence of 5 µM HspB8-HspB1 swap variants.

**Figure S5.**
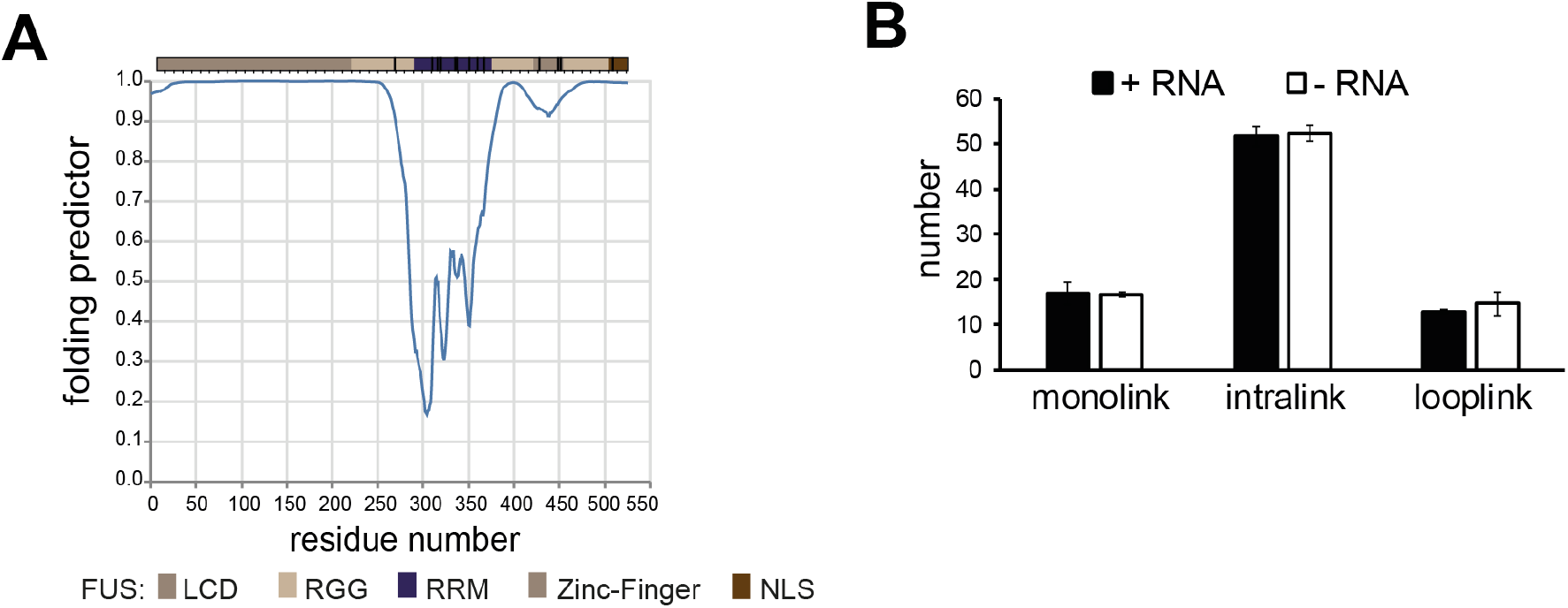
Related to Figure 5. (**A**) A disorder prediction plot (53) predicts the FUS-RRM to be folded. (**B**) FUS_m_:HspB8 condensates were crosslinked in the presence (black) or absence (white) of a customized RNA oligonucleotide previously shown to bind to FUS (40) and analysed by LC MS/MS (n=3; FDR ≤ 0.05).

**Movie 1. Time lapse of FUS fiber growth**

Time lapse movie of FUS_m_ fiber growth in the presence and absence of HspB8. The y direction images are scaled by a factor of 3 to match the scale bar for the other maximum projection. The histograms/contrast are automatically set using the enhance contrast command in FIJI with 0.3% saturated pixels.

## Acknowledgements

We thank the Protein Expression Purification and Characterization Facility, the Light Microscopy Facility and the Mass Spectrometry Facility at the Max Planck Institute of Molecular Cell Biology and Genetics in Dresden, the Bioimaging Centre (BIC) at the University of Konstanz and the Centro Interdipartimentale Grandi Strumenti (CIGS) at the University of Modena and Reggio Emilia. We thank Marit Leuschner and Anne Schwager for assistance with cell culture. We thank members of the Alberti, Hyman and Stengel Labs and Dewpoint Therapeutics for helpful discussions and critical reading of the manuscript. This work was supported by the Konstanz Research School Chemical Biology (KoRS-CB) and the Cluster of Excellence “Physics of Life”, TU Dresden, Dresden, Germany. S.A. and S.C. are grateful for the EU Joint Programme— Neurodegenerative Disease Research (JPND) project. S.C. acknowledges funding from AriSLA Foundation (Granulopathy and MLOpathy), MAECI (Dissolve_ALS) and MIUR (Departments of excellence 2018–2022; E91I18001480001). F.S. is funded by the Emmy Noether Programme of the DFG (STE 2517/1-1).

## Author Contributions

E.E.B., J.F., S.A., S.C., A.A.H. and F.S. conceived the study and experimental approach; E.E.B and M.R.-G. expressed and purified all proteins; E.E.B carried out molecular aging, partitioning and photobleaching experiments with help from X.Y. and J.W., FUS_m_ΔRRM aging assays were carried out by M.N. and J.F.; L.J. performed and analyzed time lapse microscopy experiments with help from E.E.B.; M.J. performed and analyzed fusion optical tweezer experiments with help from E.E.B.; D.M., X.Y. and L.M. carried out immunostaining and stress granule analyses; I.P. constructed all BAC cell lines; A.P. constructed all clones with help from E.E.B.; J.F. and E.E.B., prepared samples for XL-MS experiments; J.F. performed and analyzed XL-MS, q-XL-MS and aging-q-XL-MS experiments; K.M.K. wrote scripts for q-XL-MS and time-resolved-q-XL-MS analysis; E.E.B., S.A., A.A.H. and F.S. wrote the paper with input from all authors.

## Materials & Correspondence

Correspondence and material requests should be addressed to: &

## Materials & Methods

### Protein expression and purification

FUS-G156E-GFP (“FUS_m_) was purified as described (4). HspB8 and corresponding variants were subcloned in a pET11d vector as N-terminal 3C protease-cleavable GST fusion proteins. Fusion proteins were expressed and purified from BL21 Codon RIL (Stratagene). Expression was induced by adding 0.15 mM IPTG for 4,5 h at 37 °C. Bacteria were lysed in 1 x PBS, 5 mM DTT, 1 mM EDTA with EDTA-free Protease inhibitors tablet (Roche) and GST purified. Eluates were dialyzed with a 3,500 Da MWCO membrane against 1 x PBS, 5 mM DTT and cleaved with PreScission protease. Cleaved off GST was removed by reverse GST purification. HspB8 proteins were subjected to ResourceQ ion exchange chromatography, concentrated, dialyzed to HspB8 buffer (20 mM Tris, pH 7.4, 20 mM KCl, 1 mM DTT) and validated by MS.

### In vitro experiments

Frozen aliquots of FUS_m_ were thawed for 10 min at RT, cleared from aggregates by centrifugation for 1 min at 21,000 x g using a 0.2 µm spin filter device. Molecular aging experiments were performed according to (54) at 5 µM FUS_m_ in reaction buffer (20 mM Tris-HCl, pH 7.4, 75 mM KCl, 0.75 % Glycerole, 1 mM DTT). Fluorescence recovery after photobleaching (FRAP) experiments were performed and analyzed according to (4) at 5 µM FUS_m_ in reaction buffer. For partitioning experiments, HspB8 and variants thereof were labelled with Cyanine-3-monosuccinimidyl ester (AAT bioquest, ABD-141) at equimolar ratio in HspB8 buffer and excess dye was removed by dialysis against HspB8 buffer with 1 mM DTT. Labelled HspB8 was mixed with unlabelled protein at a molar ratio of 1:20 and 5 µM FUS_m_ was mixed with 5 µM total HspB8 in reaction buffer. Samples were applied into an imaging chamber with a cover slip passivated with polyethylene glycol. Fluorescence and DIC microscopy were performed on a confocal spinning disk microscope. Images were analyzed using Fiji software (55).

### Optical Tweezer experiments

To characterize the material state of FUS_m_ condensates with or without HspB8, controlled fusion experiments were performed in a custom-build dual-trap optical tweezer microscope (4, 56). 5 μM FUS_m_ condensates were phase-separated at T0 in reaction buffer with or without 20 μM HspB8 and immediately applied to a sample chamber. Two condensate droplets were trapped in two optical traps of the same trap stiffness at low overall light intensity to minimize local heating. With the first trap stationary, the second trap was moved to bring the droplets into contact and initiate coalescence, after which both traps were kept stationary. Laser signals and bright-field microscopy images were simultaneously recorded. Signals from the two traps – equal in magnitude, opposite in sign – were combined into the differential signal, from which coalescence relaxation times were deduced (22). To quantify the coalescence dynamics and account for droplets of different sizes, the relaxation time was normalised by the geometric radius of the two fusing droplets. Successful droplet coalescence was scored as yes (1) or no (0) depending on whether the process resulted in a near spherical shape of the final droplet within 60 s. This duration was an order of magnitude longer than the earliest coalescence relaxation times under all conditions. Coalescence success / failure data of FUS_m_ without HspB8 were fit with a logistic regression model to estimate the half-life of liquid-like FUS_m_ condensates.

### Time lapse microscopy of fiber growth

FUS_m_ was aged in a centrifuge tube for > 24 hours to allow most of the protein to convert to fibrous/aged material. A small amount of aged material was flowed into a custom-built flow cell which includes upper and lower glass surfaces; the bottom glass surface was passivated with polyethylene glycol. After incubation, the chamber was flushed with freshly formed FUS_m_ condensates in reaction buffer either in the presence or absence of 20 µM HspB8. An image stack representing a volume of approximate 100 µm^3^ and a voxel size of 0.1 µm x 0.1 µm x 0.3 µm was acquired every 30 minutes using a spinning disk confocal equipped with a glycerol immersion 60x objective. In the resulting image stacks, fibers tend to be relatively dim with bright cores. To produce an image which allows for good visualization of the process of fiber growth, we smoothed each stack and subsequently applied an enhanced local contrast method (CLAHE). This method uses tiles throughout the image and calculates an appropriate contrast for each tile. We used CLAHE implemented in ImageJ with a block size of 30. For each stack, we create a maximum projection of the resulting stack that produces a single image. Finally, we used an image registration method (using the stackreg plugin in ImageJ) to remove any small translational drift which occurs through the process. The resulting movie is shown as *Movie 1* in the Supplementary Data.

### Image analysis to identify droplets and fibers

The identification of fibers and droplets in images was carried out using custom-made scripts in MATLAB. In short, each image is resized 8 times using a bicubic interpolation. An image is subsequently automatically thresholded using Otsu’s method through the imthresh command. Objects are identified as regions of connected pixels. Any objects which intersect the picture border or are very small are discarded from further analysis. Fibrous objects are identified as either objects with an eccentricity above 0.7 or, if the eccentricity is low, as objects that have a roughness above 6.4 pixels. The eccentricity is found by fitting an ellipse to a connected region and is defined as the ratio of the distance between the foci of the ellipse and its major axis length (implemented using the eccentricity argument in the regionprops command). To determine the roughness, each object is fit by a circle. The roughness is defined as the mean distance between the object border and the circular fit. Objects with low surface roughness as well as low eccentricity were considered droplets. All objects and their subsequent classification are also reviewed finally by eye to ensure that the parameters for the images are set properly.

### Immunostaining of sHSPs in stressed cells

HeLa Kyoto wildtype and HeLa Kyoto FUS-GFP BAC cells were cultured in Dulbecco’s Modified Eagle Medium (DMEM) containing 4.5 g/l glucose (Gibco Life Technologies) supplemented with 10% fetal bovine serum, 100 U/ml Penicillin + 100 μg/ml Streptomycin. 250 μg/ml Geneticin (all Gibco Life Technologies) was added to the HeLa Kyoto FUS-GFP BAC cells. Cells were maintained at 37°C in a 5 % CO2 incubator (Thermo Fisher Scientific). HeLa Kyoto FUS-GFP BAC cells were described previously (12). For immunostaining of HspB8-WT and HspB8-0R in stress granules, HeLa Kyoto cells were transfected with 200 ng of plasmids coding for HspB8-0R-3xmyc or HSPB8-WT-3xmyc using Lipofectamine 2000 (Life Technologies) following manufacturer instructions. 24 hrs post-transfection cells were subjected to heat shock in a water bath at 43.5 C for 1 hr. Cells were fixed with 3.7% formaldehyde for 9 min at RT and permeabilized with acetone for 5 min at −20C and stained with c-Myc (9E10, Santa Cruz Biotechnology) and eIF4G (H-300, Santa Cruz Biotechnology) specific antibodies. Secondary antibodies used were anti-mouse Alexa Fluor 594 (A-21203, Thermo Scientific) and anti-rabbit Alexa Fluor 488 (A-21206, Thermo Scientific).

### Crosslinking of molecular condensates

Frozen aliquots of FUS_m_ and FUS_m__K9 protein were thawed for 10 min at RT, cleared from aggregates by centrifugation for 1 min at 21,000 x g using a 0.2 µm spin filter device and subsequently diluted in water to a low salt solution (final concentration of 75 mM KCl) to induce phase separation or into a high salt solution (final concentration of 500 mM KCl) to prevent phase separation. In order to reconstitute FUS_m_:HspB8 condensates HspB8-WT or HspB8-K141E mutant were added to FUS_m_ condensates at equal mass ratio and subsequently incubated on ice for 10 min to allow for sufficient mixing. All molecular condensates were crosslinked by addition of H12/D12 DSS (Creative Molecules) at a molar ratio crosslinker to lysines of ∼2.7 for 30 min (except for the time-course experiments, see below) at 37°C shaking at 650 rpm in a Thermomixer (Eppendorf). Protein samples were quenched by addition of ammonium bicarbonate to a final concentration of 50 mM and either directly evaporated to dryness or after an additional centrifugation step for 60 min at 21 000 x g in order to separate the condensed from the dilute phase. The dilute phase containing supernatant was transferred to a fresh tube and both phases were evaporated to dryness.

### Crosslinking coupled to Mass Spectrometry (XL-MS)

Crosslinked samples were processed essentially as described (57). In short, the dried protein samples were denatured in 8 M Urea, reduced by addition of 2.5 mM TCEP at 37°C for 30 min and subsequently alkylated using 5 mM Iodacetamid at RT for 30 min in the dark. Samples were digested by addition of 2 % (w/w) trypsin (Promega) over night at 37°C after adding 50 mM ammonium hydrogen carbonate to a final concentration of 1 M urea. Digested peptides were separated from the solution and retained by a C18 solid phase extraction system (SepPak Vac 1cc tC18 (50 mg cartridges, Waters) and eluted in 50 % ACN, 0.1 % FA. After desalting the peptides were evaporated to dryness and stored at −20°C. Dried peptides were reconstituted in 30 % ACN, 0.1 % TFA and then separated by size exclusion chromatography on a Superdex 30 increase 3.2/300 (GE Life Science) to enrich for crosslinked peptides.

The three early-eluting fractions were collected for MS measurement, evaporated to dryness and reconstituted in 5 % ACN, 0.1 % FA. Concentrations were normalized by A215 nm measurement to ensure equal amounts of dilute and condensed phase and peptides separated on a PepMap C18 2µM, 50 µM x 150 mm (Thermo Fisher) using a gradient of 5 to 35 % ACN for 45 min. MS measurement was performed on an Orbitrap Fusion Tribrid mass spectrometer (Thermo Scientific) in data dependent acquisition mode with a cycle time of 3 s. The full scan was done in the Orbitrap with a resolution of 120000, a scan range of 400-1500 m/z, AGC Target 2.0e5 and injection time of 50 ms. Monoisotopic precursor selection and dynamic exclusion was used for precursor selection. Only precursor charge states of 3-8 were selected for fragmentation by collision-induced dissociation (CID) using 35 % activation energy. MS2 was carried out in the ion trap in normal scan range mode, AGC target 1.0e4 and injection time of 35 ms. Data were searched using *xQuest* in ion-tag mode. Carbamidomethylation (+57.021 Da) was used as a static modification for cysteine. As database the sequences of the measured recombinant proteins and reversed and shuffled sequences were used for the FDR calculation by *xProphet*.

Experiments were carried out in three biologically independent sets of experiments (meaning separate batches of expressed protein). For one set of experiments each sample was independently crosslinked in triplicates and each of these was measured in technical duplicates. Crosslinks were only considered, if they were identified in two out of three replicates with a deltaS < 0.95, a minimum Id score ≥ 20 and an ld score ≥ 25 in at least one replicate (filtering was done on the level of the unique crosslinking site) and an FDR ≤ 0.05 as calculated by *xProphet* for at least one replicate.

### Quantitation of Crosslinked Peptides from Condensates (qXL-MS)

#### Quantitation

Initial processing of identified crosslinked peptides for quantitation was performed essentially as described (31). In short, the chromatographic peaks of identified crosslinks were integrated and summed up over different peak groups for quantification by *xTract* (taking different charge states and different unique crosslinked peptides for each unique crosslinking site into account). Only high-confidence crosslinks that fulfilled the above introduced criteria were selected for further quantitative analysis.

The resulting *bagcontainer*.*details*.*stats*.*xls* file was used as an input for in-house scripts developed for this manuscript. The bag container contains all experimental observations on a peptide level as extracted by *xTract* (e.g. peptide mass, charge state, the extracted MS1 peak area and any violations assigned by *xTract)*. Missing observations were replaced by imputation with random values drawn from a normal distribution based on our experimental distribution. Here, the log-normal experimental distribution of measured MS1 peak areas was converted to a normal distribution by log2-conversion. Of the resulting normal distribution, the mean and standard deviations were determined. The mean was shifted downward while the width was decreased in order to obtain the distribution to draw imputed values from, following the same procedure and parameters as described for Perseus (58) (width: 0.3 and down shift: 1.8).

Data were additionally filtered using a light-heavy filter as described (59) and peptides with a light-heavy log2ratio <-1 or >1 were excluded from further analysis. Experiments were normalized by their mean MS1 peak area using the mean of all experiments as reference. The ratio of each experiment compared to the reference was computed and all observed MS1 areas were multiplied by this experiment-specific ratio to receive the same mean for all experiments. In addition, replicates were normalized within each experiment. Thus, the mean of each biological and technical replicate within an experiment is shifted to the mean of an experiment in the same way as described above.

In a next step log2ratios were calculated as the difference between the log2-converted MS1 peak areas (instead of the ratio). Here, the MS1 area for each experiment was shifted into a log2 scale after all summing operations but before taking any means, allowing us to calculate meaningful standard deviations between biological replicates and to avoid the influence of outliers in the original log-normal scale. P-value calculations were otherwise performed as described (59), with one notable exception: MS1 peak areas were not split by technical replicates in order to avoid artificially improved p-values with increasing numbers of technical replicates. FDR values were p-values corrected for multiple testing, following the Benjamini–Hochberg procedure.

#### Significance

Only high-confidence crosslinking sites (see above) that were detected reliably and consistently with a deltaS < 0.95, a minimum Id score ≥ 20 and a ld score ≥ 25 in at least one replicate (filtering was done on the level of the unique crosslinking site) and an FDR ≤ 0.05 were used for quantitation. Changes in crosslinking abundances were throughout the paper quantified against the dilute phase (i.e. relative enrichment within droplets is shown in green; relative decrease in red). Only crosslinking sites that were up – or downregulated two-fold or more (log2ratio ≥ 1 or ≤ −1 and FDR ≤ 0.05) in at least two biological replicate sets of experiments and in addition contained no opposing regulation in any replicate set were considered significant.

### Time-resolved quantitative crosslinking coupled to mass spectrometry

Fresh FUS_m_ condensates formed under low salt (75 mM NaCl) conditions were left shaking at 650 rpm in a Thermomixer (Eppendorf) at 28°C and monitored by fluorescence microscopy at regular intervals until conversion into fibers. The stock solution was aliquoted prior to dilution into low salt buffer to induce condensation and aliquots (n=3) were crosslinked for 5 minutes and flash-frozen in liquid nitrogen at indicated timepoints: T1 to T6 (0 hrs, 20 min, 40 min, 1hrs, 1hrs 20 min, 1hrs 40 min; condensates), T7-T9 (2 hrs 20 min, 2 hrs 40 min, 3 hrs; gels) to T10-T11 (12 hrs and

24 hrs; fibers). While thawing, 1M ammonium bicarbonate was added to a final concentration of 50 mM and samples were evaporated to dryness. Crosslinks were subsequently identified and quantified exactly as described above.

### Visualization of crosslink data

In order to both validate and visualize the crosslink information multiple in-house scripts have been written. The visualization scripts either interface directly with the quantitation script described above or use *xTract*-like output for an input. In either case, the filtering and significance criteria as described above for the quantitation script were used. Crosslink data were transformed via pandas (version 1.0.3) for assessment. Figures 1B, 5B, S3F and S6 were created with altair (version 4.1.0) running on Python version 3.7.6. Figures 1C and S1A (left and right Panel) were created with seaborn (version 0.9.0) running on python (version 3.7.2). Figure S1A (middle panel) was created using the in-built pandas dataframe plot functions.

### RNA competition assay

FUS_m_:HspB8 condensates were prepared as described and either incubated with RNA oligonucleotide PrD (40) in sub-stoichiometric amounts (3 times molar excess of FUS_m_) or equal volume of water. Samples were checked by microscopy before crosslinking to ensure that addition of RNA did not dissolve the condensates. The condensed phase was separated from the dilute phase by centrifugation and the concentration normalized prior to MS measurement as described above.

## Data availability

All data generated or analysed during this study are included in this published article (and its supplementary information files). The MS data (raw files, *xQuest, xTract* and in-house quantitation output files) have been deposited to the ProteomeXchange Consortium via the PRIDE (60) partner repository with the dataset identifier PXD021114 (Username; Password: 5atfkbP8) and PXD021115 (Username; Password: UZW7Gnr5).

